# A Bioprinted Head and Neck Cancer Organoid-Based Platform for Evaluating Multimodal Therapies

**DOI:** 10.64898/2026.05.20.726741

**Authors:** Luda Lin, Krishna K. Bommakanti, Christian Wooten, Alfredo Enrique Gonzalez, Yazeed Alhiyari, Jonathan Levi, Bowen Wang, Andreanne Sannajust, Lauran K. Evans, Peyton Tebon, Maie A. St. John, Alice Soragni

**Author notes:** Correspondence to AS and MSJ.

## Abstract

Treatment of advanced head and neck squamous cell carcinoma (HNSCC) often involves radiotherapy combined with chemotherapy, targeted therapy, or immunotherapy. However, due to its anatomical and molecular heterogeneity, identifying the most effective treatment for each patient remains a major clinical challenge. To address this need, we developed a high-throughput organoid-based drug screening platform that uses patient-derived organoids to assess candidate treatment regimens. We validated the platform by establishing bioprinted 3D organoids of human HNSCC cell lines and exposing them to X-ray radiation in combination with various small-molecule drugs and biologics. We quantified viability using ATP release assays and assessed extracellular matrix (ECM) invasion with a machine learning-based brightfield image analysis pipeline. Proof-of-concept experiments with HPV-negative HNSCC lines (HN30 and HN31, established from primary and metastatic disease from the same patient) and HPV-positive HNSCC cells (SCC154) revealed different therapy agents that can radiosensitize each cell line. Image analysis showed that copanlisib, afatinib, and ibrutinib could limit ECM invasion of HN31, while the AKT inhibitor ipatasertib promotes invasion of HN30 cells, consistent with previous studies. Application of the platform to patient-derived HPV+ oropharyngeal tumor organoids showed that they shared sensitivity to several agents while also exhibiting differences against certain therapies. Cetuximab, sorafenib, and nedisertib significantly radiosensitized organoids from two clinical samples. This work demonstrates the feasibility of performing sensitivity screening by integrating bioprinting, conventional viability assays, and advanced image analysis techniques. This platform has the potential to enable a personalized therapeutic pipeline for patients with advanced HNSCC, optimizing responses to radiotherapy and targeted agents to improve clinical outcomes while avoiding modulators that may promote tumor invasion.

## Introduction

Head and neck cancer (HNC) is the sixth most prevalent cancer in the world, with over one million new cases per year [1]. More than 90% of these malignancies are classified as head and neck squamous cell carcinomas (HNSCC), which are conventionally related to either tobacco and alcohol use [2], or viral infections (i.e. human papillomavirus (HPV) in US [3–6] and Epstein-Barr virus (EBV) in Southern and Southeastern Asia [7,8]). In addition to being anatomically heterogenous (e.g. originating in the oral cavity, larynx, and oropharynx), the genetic comparison between non-HPV-induced (HPV-) and HPV-induced (HPV+) subtypes revealed the presence of distinct sets of mutations responsible for malignant transformation [9,10].

Different anatomical locations, HPV status, and mutations cause major differences in HNC tumor physiology and microenvironment, resulting in high variability of patient response to treatment. This limits the effectiveness of available therapies, including chemotherapy, targeted drugs and immunotherapies [11]. This problem highlights the need for a personalized therapeutic pipeline for HNSCC patients, especially patients with advanced disease that cannot be managed using the typical standard of care (surgery, followed by radiotherapy and, in more advanced cases, chemotherapy). Lymph node metastasis is another major problem as it occurs in a large proportion of patients (25– 45% of all HNC cases) and leads to a significantly worse prognosis [12,13]. Furthermore, distant metastasis, found in about 10% of HNC cases, leads to a median overall survival of 10 months and a 5-year survival rate of 9% [9,14,15]. Therefore, finding therapeutic agents and regimens that can inhibit cancer invasion and metastasis in HNC patients remains a critical medical need [16,17]. However, typical 2D cell-based drug screening strategies that are used to discover therapeutics with anti-cancer activities are poor models of cancers with high heterogeneity and metastatic cancers.

Over the last 15 years, organoids, 3D multicellular bodies grown from cell lines or patient clinical samples embedded in a hydrogel, have emerged as more physiologically relevant models of cancer as they better recapitulate complicated structures and functions of in vivo tissues when compared to traditional cell lines [18–20]. Patient-derived organoids have been used to screen chemotherapies, targeted agents and radiotherapies in HNCs [21–27], as well as to discover invasion signatures of HNCs [27], evaluate effectiveness of EGFR-targeted photodynamic therapy [29], and identify subtype-specific therapeutic options in nasopharyngeal carcinoma [30]. Although these studies represent important advances, current patient-derived organoids have limitations, most notably their mechanical fragility necessitating gentle handling conditions. As such, these organoids remain incompatible with automation, a necessary step for improving throughput and translating organoid-based screens into a clinical-grade personalized medicine platform.

To address these challenges, we developed an efficient ring-like organoid-based drug screening platform, which protects organoids from disruption or mechanical forces and is compatible with automated liquid handlers [28,29] and an automated process of organoid seeding that uses a bioprinter [30]. This platform is similar to organoid systems we developed previously to reveal drug sensitivity and provide therapy recommendations for several rare and heterogeneous tumors [31–33]. The new HNSCC platform uses an endpoint ATP assay to assess cell viability. We used it to screen a small library of 33 cancer drugs including chemotherapy agents, targeted therapies, and immunotherapies, and to map how different subtypes of HNSCC (HPV− and HPV+) respond to treatment. We also examined how co-treating with radiation and the therapeutic agent affects viability, especially focusing on the discovery of drugs that function as radiosensitizers. Furthermore, we showed that the platform is fully compatible with label-free brightfield image analysis with organoid segmentation. When combined with machine learning (ML)–based image analysis, our platform enabled the measurement of size and shape of organoids, and how these characteristics change upon treatment. Moreover, we showed that our organoid platform is compatible with immunofluorescence imaging, providing a more granular view of cellular perturbations caused by different treatments. When applied to analyze clinical samples, the platform identified actionable opportunities, including drugs that showed high efficacy across organoids derived from four patients. Overall, this patient-derived organoid platform represents a significant step towards a personalized therapeutic pipeline for patients with advanced HNSCC, maximizing responses to radiotherapy and targeted agents to improve outcomes.

## Results

### A Proof-of-Concept Platform Captures Different Sensitivity of Organoids Derived from Different HNSCC Cell Lines

To prototype our HNSCC organoid-based platform and assess its capacity to detect clinically relevant differences across HNSCC subtypes and stages, we generated organoids from three well-established HNSCC cell lines: HN30, HN31, and SCC154. In brief, HN30 and HN31 are isogenic cell lines derived from a primary tumor and lymph node metastasis sample, respectively, from the same HPV-HNSCC patient, while SCC154 originated from a different patient whose disease was HPV+. These cell lines also have a different TP53 mutational status (wild-type TP53 in HN30 and SCC154, and mutant TP53 in HN31) [34]. As such, organoids derived from these cell lines provide distinct models of human HNSCC, enabling us to test the performance of our platform across three clinically relevant conditions (HPV+ vs. HPV-, and primary vs. metastatic tumor subtypes). The organoid platform was prepared as described in the Methods section, using previously described procedures [28–33].

To assess the platform’s performance, we first administered a single dose of radiation ranging from 2 – 16 gray (Gy) to organoids derived from HN30, HN31, and SCC154 to measure their radiosensitivity. We observed that while HPV+ SCC154 organoids were sensitive to radiation (less than 40% viability at 16 Gy; **Figure 1A**), viability of HPV-HN30 and HN31 was affected to lesser extent (82.9% and 78.6% viability at 16 Gy, respectively; **Figure 1A**), which agrees with previously reported differences in radiosensitivity between HPV+ and HPV-HNSCC cell lines [35,36]. According to the results of radiation gradient assays, we chose 2 Gy radiation for SCC154, and 4 Gy radiation for HN30 and HN31 organoids as conditions for our follow-up experiments.

**Figure 1.**
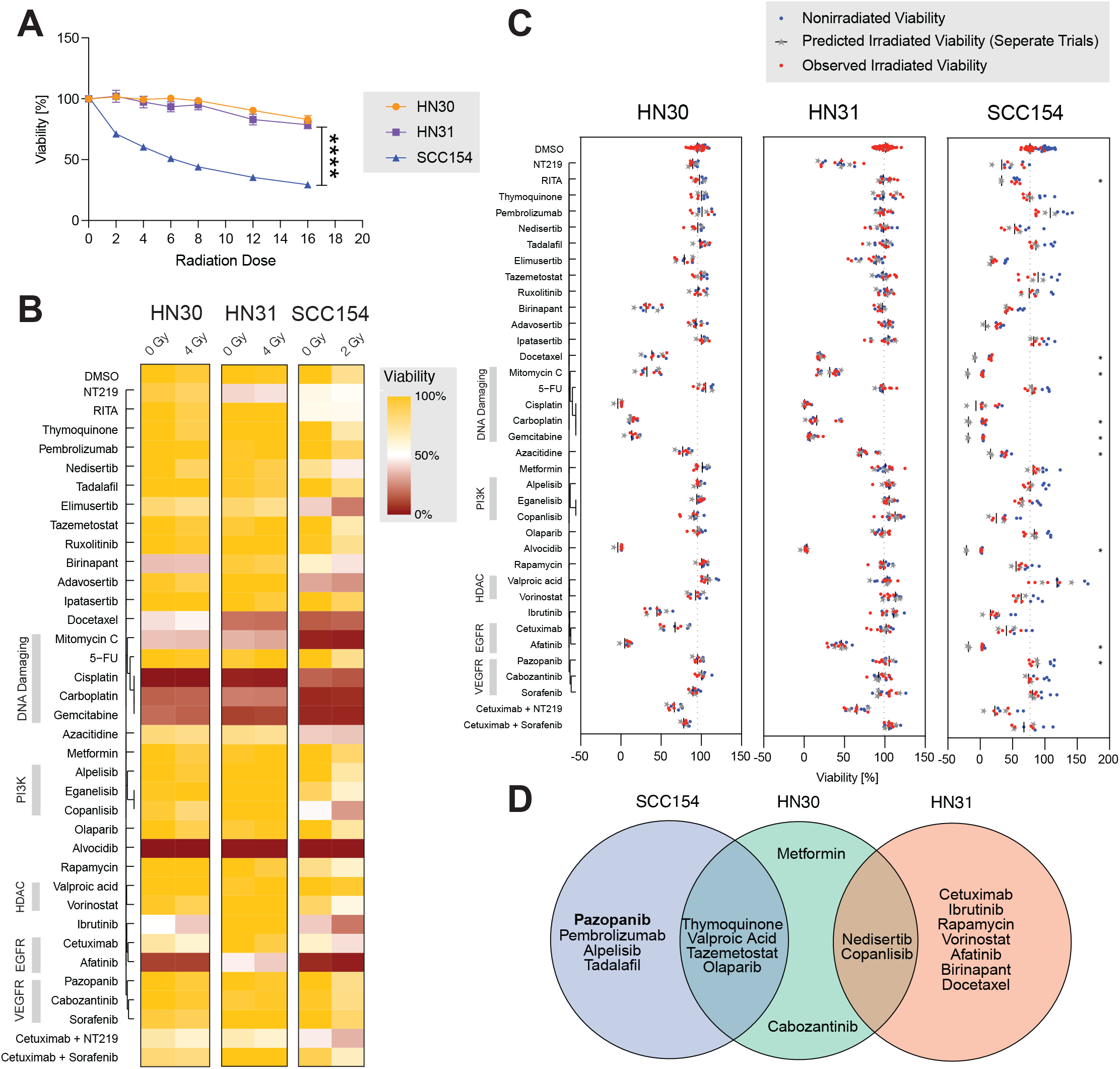
Assessment of HN30, HN31 and SCC154 organoid-based assays. **(A)** Dose-response curves of organoid viability of HN30, HN31 and SCC154 exposed to radiation of different intensities in the radiation gradient assay. Data are represented as mean ± SEM. ****: q < 0.00001. (**B**) Heatmap of high-throughput screening results of HN30, HN31, and SCC154 organoids for 33 therapy agents and 2 combinations with cetuximab, with or without radiation of specific doses for each cell line. The color represents the normalized cell viability %. The drugs are clustered using the Jaccard distance based on common protein targets. (**C**) Scatter plots of high-throughput screening results for HN30, HN31, and SCC154 cell line–derived organoids assessing the combination effects of therapeutic agents with radiation. Blue dots represent the normalized cell viability % of individual wells without radiation. Grey stars represent predicted post-radiation viability, calculated with a linear interaction model based on the agent-specific effect, derived from the average viability of all nonirradiated wells within the same trial. Black vertical bars represent mean predicted viability across trials. Red dots represent observed viability % of wells treated with both the therapeutic agent and radiation. Synergistic effects are shown if red dots lie to the left of the black bar, which indicates lower-than-expected viability. Student’s t-test was used to compare predicted and observed irradiated values. *: p < 0.05. (**D**) The Venn diagram denoting radiosensitizer candidates of HN30, HN31 and SCC154. Bold indicates a significant radiosensitizing effect, defined as p < 0.05 for predicted vs. observed irradiated values shown in Figure 1C. See also **Supplementary Figures 1-3**.

Next, we performed an unbiased drug screening assay of our organoids using a small library of 33 anticancer drugs (**Supplementary Table 1**), including chemotherapy agents (e.g. DNA damaging drugs such as cisplatin), targeted therapies (e.g. inhibitors of EGFR, PI3K, HDAC, VEGFR, and PARP), and immunotherapy drugs (**Figure 1B**). Several drugs, including chemotherapy agents cisplatin, carboplatin, gemcitabine, mitomycin and docetaxel, pan-CDK inhibitor alvocidib, and EGFR inhibitor afatinib, were effective against all three organoids, leading to a viability lower than 50% (**Figure 1B**). Other drugs including Wee-1 inhibitor adavosertib, ATR inhibitor elimusertib, DNA methylation inhibitor azacitidine, pan-PI3K inhibitor copanlisib, and MDM2 inhibitor RITA were significantly more effective against SCC154 organoids than the two derived from HPV-cell lines. A comparison of drug sensitivities between HN30 and HN31 organoids (**Supplementary Figure 1**) showed that IAP inhibitor birinapant, EGFR inhibitor afatinib, and the combination of EGFR inhibitor cetuximab and sorafenib are significantly more effective against HN30, whereas IRS and STAT3 inhibitor NT219 and mitotic inhibitor docetaxel were more effective against HN31. The combinations of cetuximab with NT219 and cetuximab with sorafenib showed significant antagonist effects (**Supplementary Figures 2-3**): cetuximab with NT219 in HN31 (6 estimate 0.2502, p-value 0.0004) and SCC154 (β6 estimate 0.2563, p-value 0.0156), and cetuximab with sorafenib in HN30 (β6 estimate 0.1627, p-value 0.06022) and SCC154 (β6 estimate 0.2292, p-value 0.0228).

To examine whether our organoid-based assays could identify radiosensitizers, defined as agents that synergize with radiation, we applied a linear interaction model comparing the predicted viability of organoids treated with radiation or individual drug agents alone to the predicted viability observed following combined radiation and drug treatment (**Figure 1C**). While most significant combination effects were antagonistic, with observed viability exceeding predicted viability, pazopanib, a multi-kinase inhibitor, significantly radiosensitized SCC154 cells, resulting in an observed post-irradiation viability of 78.2% compared to a predicted viability of 88.4% (p = 0.0023) (**Figure 1C-D**). In addition, we identified 18 other agents showing more modest trends toward radiosensitization across the three cell lines analyzed (**Figure 1D**). HN31 and SCC154 exhibited distinct radiosensitization profiles, with 9 and 8 agents identified, respectively. HN30 shared 2 agents with HN31 and 4 with SCC154, while 2 additional agents were unique to HN30 (**Figure 1D**). These similarities and differences in treatment response could reflect shared molecular and biological features of the parental cells. For example, HN30 and HN31 are both HPV-negative, HN30 and SCC154 are derived from primary tumors, while HN31 is the only metastatic, TP53-mutant cell line in the cohort. However, the specific pathways and mechanisms underlying these response patterns require further investigation.

Overall, assessment results suggest that our HNSCC organoid platform offers a viable strategy for measuring radiosensitivity and drug response in a high-throughput manner compatible with screening patient-derived organoids to identify the most effective treatment regimens.

### Machine Learning-Based Image Analysis of Organoids Reveals Influence of Therapy Agents and Radiation on Organoid Size and Invasion Ability

Although the ATP release assay is considered the gold standard for measuring cell viability, it is typically used as a single-timepoint viability readout, i.e., at the end of the incubation period. This means that the assay cannot visualize and detect changes that may occur during the incubation period. To better evaluate the effects of the administered treatments on the organoids throughout the incubation period, we combined daily organoid imaging with a ML–based image analysis workflow to measure their size and morphology in brightfield images. This approach allowed us to track how growth and invasion of organoids change in response to administration of drugs or radiation (**Figure 2A-C, Supplementary Figures 4-5**). We observed that untreated HN31 organoids grew the fastest (normalized growth = 10.14), followed by HN30 (normalized growth = 6.66) and SCC154 (normalized growth = 3.37). Many HN31 organoids were invasive, with sprouts growing over time into the adjacent ECM and leading to a spiky or stellate morphology (low circularity value of 0.58 on Day 9; **Figure 2B**), whereas HN30 and SCC154 organoids were generally spheroidic (circularity values 0.66 (**Figure 2A**) and 0.69 (**Figure 2C**) on Day 9, respectively). We also noted that organoids responded to different drugs differently; for example, ibrutinib and adavosertib caused HN30 (**Figure 2A**) and SCC154 organoids (**Figure 2C**) to decrease in size respectively, consistent with their toxicity on the corresponding cell lines (**Figure 1B**). Ibrutinib and cetuximab led HN31 organoids to lose invading sprouts and become rounded (**Figure 2A-B**), whereas ipatasertib led to a slight size increase and a spikier shape of HN30 organoids (**Figure 2A**). This suggests that monitoring how physical characteristics of organoids change in response to treatment may be used for building models to predict the efficacy of HNSCC therapies, as previously proposed in other contexts [37,38].

**Figure 2.**
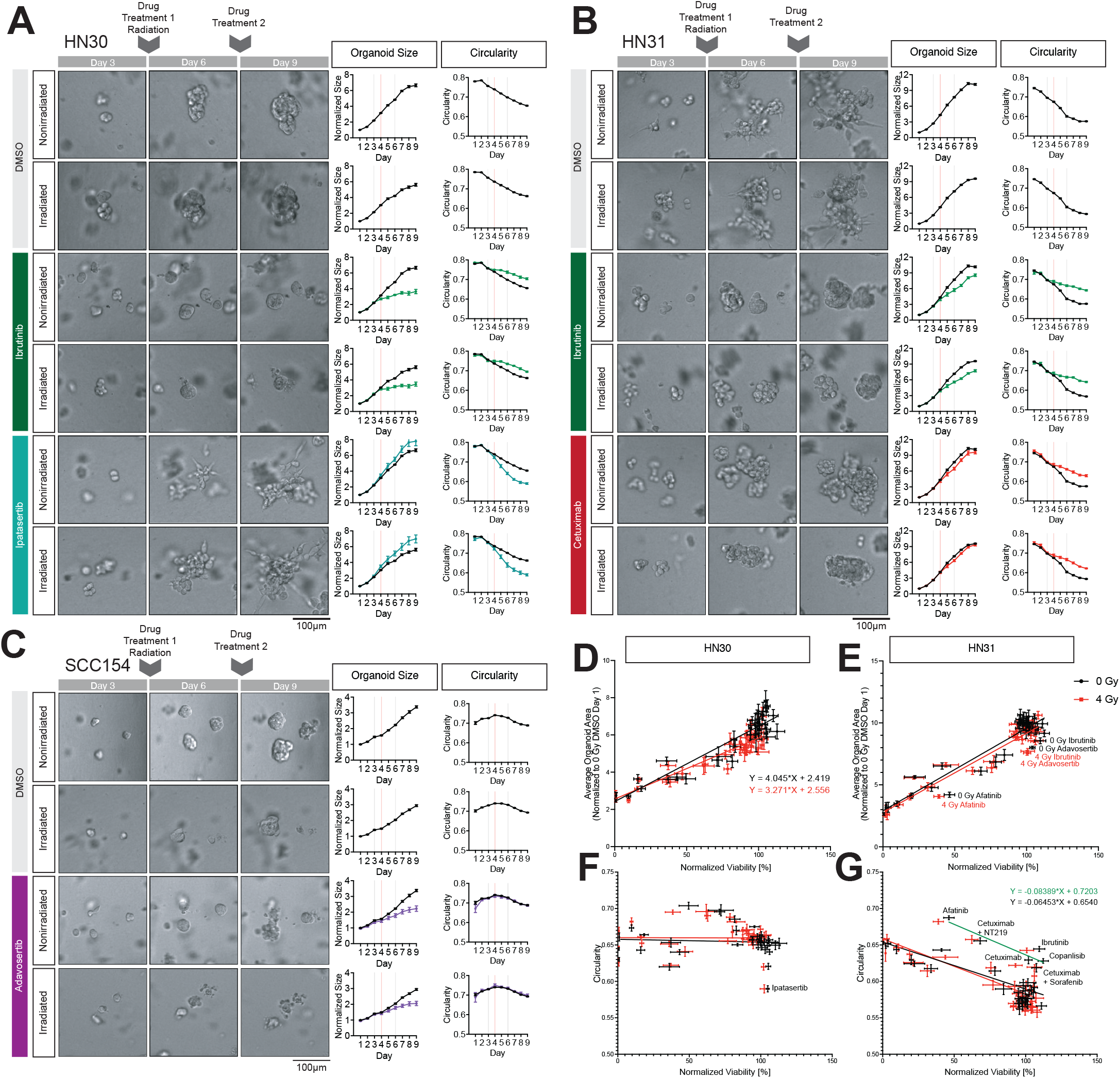
Image Analysis Results of HN30, HN31 and SCC154. **(A-C)** Representative brightfield images of organoids of HN30 **(A)**, HN31 **(B)** and SCC154 **(C)** in culture on Day 3 (Column 1), Day 6 (Column 2) and Day 9 (Column 3). Organoids were treated with therapy agents on Day 3 and Day 6, and radiation on Day 4, after imaging. Growth (Column 4) and changes of morphology of organoids (Column 5) were tracked over time by segmenting in-focus organoids in the brightfield images using a machine learning–based pipeline. Growth was measured by normalizing organoid size daily, calculated as the average cross-sectional area of all organoids across all wells within the same treatment, cell line, and trial, relative to DMSO without radiation, measured on Day 1 of culture. Data of control organoids are denoted with black lines. Data of treated organoids are denoted with respective colors. Data are represented as mean ± SEM. Morphology was measured by calculating the average circularity of all organoids across all wells of the same treatment and cell line within each trial. Scale bars: 100 μm for brightfield pictures. **(D-E)** Scatter plots of normalized viability % versus Day 9 normalized organoid size for HN31 (D) and HN30 (E). Each dot denotes one therapy agent or combination. Data without radiation are denoted with black and data with radiation are denoted with red. Data are represented as mean ± SD. Linear regression analysis was done for nonirradiated and irradiated data separately, with the black line the linear regression line of nonirradiated data and the red line that of irradiated data. **(F-G)** Scatter plots of normalized viability versus Day 9 circularity for HN31 **(F)** and HN30 **(G)**. Data are represented as mean ± SEM. Linear regression analysis was done for nonirradiated and irradiated data separately, with the black line representing the linear regression line of nonirradiated data and the red line that of irradiated data. In F, linear regression analysis was also done specifically for nonirradiated data of invasion inhibitors, with the linear regression line shown as green. See also **Supplementary Figures 4-6**.

To decouple the effects of therapy on organoid size and circularity from its toxicity, we performed a linear regression assay between normalized viability and organoid size or circularity on Day 9 and compared the predicted values with the measured organoid size and circularity (**Figure 2D-G, Supplementary Figure 5B-C**). In the case of organoid size, we noticed that adavosertib (normalized growth = 8.01, q-value = 0.000025), afatinib (normalized growth = 4.20, q-value = 0.004) and ibrutinib (normalized growth = 8.56, q-value = 0.01) significantly reduced the growth of HN31 organoids (**Figure 2E, Supplementary Figure 5**). Such significant toxicity-independent organoid shrinking effects were not found in HN30 or SCC154 organoids (**Figure 2D, Supplementary Figures 5B** and **6A**). Interestingly, the circularity of HN31 organoids was inversely correlated with viability (**Figure 2G**), potentially due to the more invasive nature of these organoids. Furthermore, four drugs and two drug combinations (afatinib, ibrutinib, copanlisib, cetuximab, cetuximab + NT219, and cetuximab + sorafenib) were notably distinguished from the data points corresponding to other treatments (**Figure 2G, Supplementary Figure 6A**). We speculate that this is due to the anti-invasion effects of those therapy agents to inhibit molecules involved in signaling pathways relevant to cancer invasion, such as EGFR and PI3K [39–45]. Ibrutinib is also reported to inhibit the invasion and migration of several solid tumors [46]. In contrast to HN31 organoids, the circularity of HN30 and SCC154 organoids did not appear to be associated with their viability since organoids maintained a spheroidal shape at all times (**Figure 2F, Supplementary Figures 5C** and **6A**). Surprisingly, we noticed that the ipatasertib treatment of HN30 organoids resulted in significantly reduced circularity. Previous studies showed that AKT1 inhibition could promote the invasion of several solid tumors, including HNSCC [47,48]. By contrast, no therapy was found to promote a reduction in circularity of SCC154 organoids (**Supplementary Figure 5C**). We also analyzed the influence of radiation on organoid size and circularity and observed that HN30 organoids treated with 4 Gy radiation (normalized size = 5.62) are smaller than those not exposed to radiation (normalized size = 6.66) (**Figure 2D**), which is not seen in HN31 or SCC154 organoids. Overall, radiation had little to no effect on how drug treatment affected organoid size or circularity (**Supplementary Figure 6**), although it did restrain overgrowth caused by metformin in HN30 and HN31, and RITA, 5-FU and tadalafil in HN31 (**Supplementary Figures 4-6**), and apparently promoted the reduction of sizes of organoids treated with sorafenib or thymoquinone (**Supplementary Figure 6**).

Taken together, monitoring organoid size and circularity provides deeper insights than the measurement of viability alone. Our data indicate that the organoid-based platform we developed is compatible with brightfield imaging that, when combined with ML–based image analysis, facilitates characterization of how specific treatment regimens affect HNSCC organoid size and shape.

### Ibrutinib has Distinct Effects on HN30 and HN31 Likely Due to its Off-Target Activity

Based on the results from our HNSCC organoid-based platform assays, we identified ibrutinib, a Bruton’s tyrosine kinase (BTK) inhibitor [49] and an approved drug for blood cancers [50,51], as an agent of interest given its unique behavior. Ibrutinib exhibited distinct effects across the organoid models, including substantially greater toxicity in HN30 organoids compared to HN31 organoids (**Figure 1B, Supplementary Figure 1**), a trend toward radiosensitization in HN31 (**Figure 1C-D**), and inhibition of invasive behavior in HN31 organoids (**Figure 2B, G**). Given that various pre-clinical trials showed that Ibrutinib was effective against multiple solid tumors including HNSCC [46], we decided to investigate the effects of ibrutinib against HN30 and HN31 organoids and whether they are mediated through inhibition of the BTK receptor.

We treated HN30 and HN31 organoids with ibrutinib and two next-generation BTK inhibitors, spebrutinib and acalabrutinib (concentrations ranging from 0.2 to 20 µM, with or without 4 Gy radiation). We found that while ibrutinib is more effective against HN30 (IC50 = 0.76 µM) than HN31 (IC50 = 3.8 µM), spebrutinib is equally effective against both (IC50 (HN30) = 9.25 µM, and IC50 (HN31) = 10.33 µM), while acalabrutinib is not toxic at concentrations as high as 20 µM (**Figure 3A**). Notably, ibrutinib appears to act as a radio-sensitizer for both HN30 (IC50 = 0.54 µM at 4 Gy radiation) and HN31 (IC50 = 3.1 µM at 4 Gy radiation), whereas spebrutinib does not show a similar synergistic effect with radiation.

**Figure 3.**
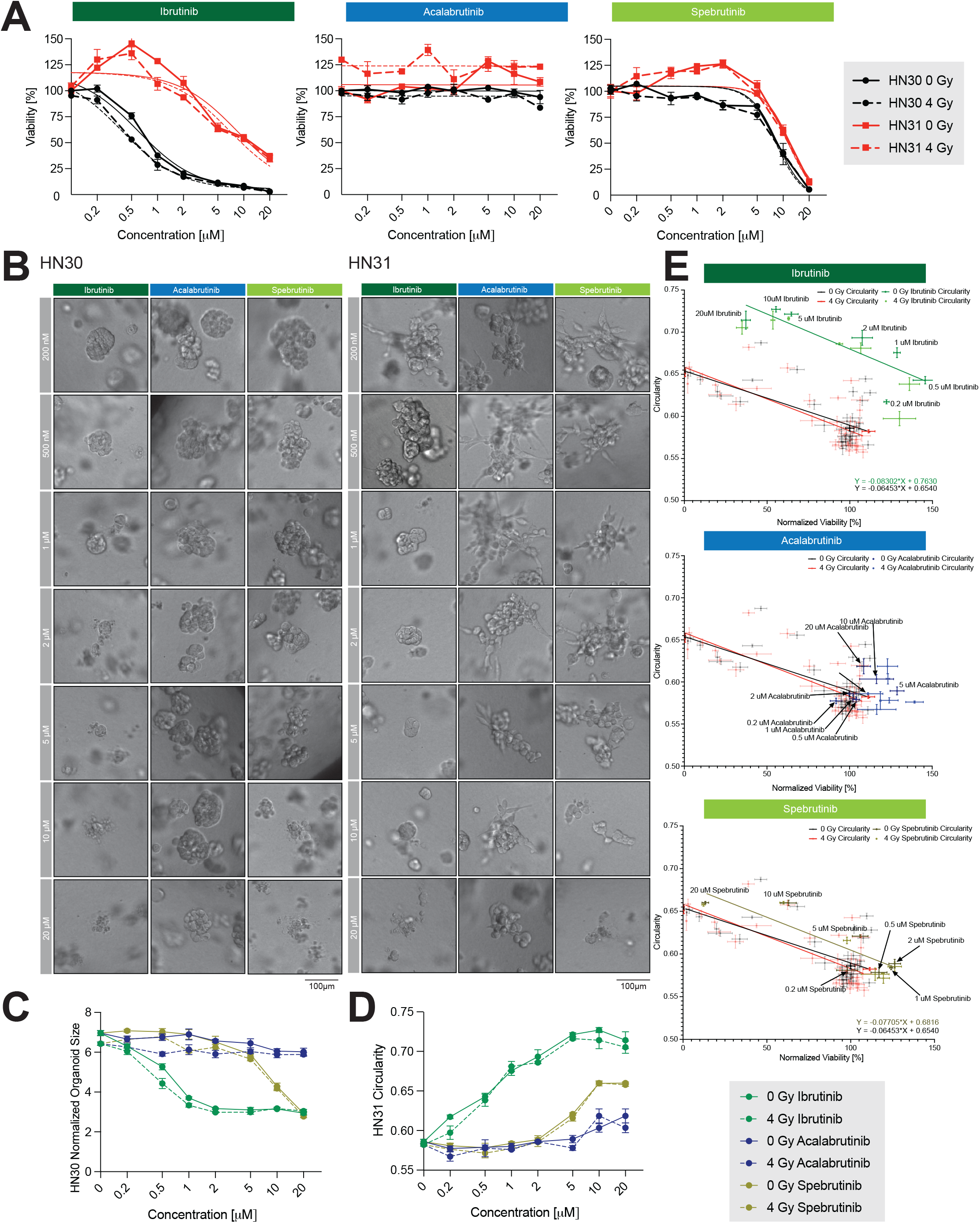
Effect of a BTK Inhibitor Ibrutinib on HN30 and HN31 organoids. **(A)** Dose-response curves of organoid viability of HN30, HN31 and SCC154 under treatment of ibrutinib, acalabrutinib or spebrutinib of different concentrations, with or without radiation. Data of HN30 organoids are shown in black, and data of HN31 are shown in red. Data of nonirradiated organoids are shown as solid lines, and data of irradiated ones as dashed lines. Line charts are overlapped with IC50 curves of each treatment, with or without radiation. Data are represented as mean ± SEM. **(B)** Representative brightfield images of HN31 organoids under ibrutinib, acalabrutinib or spebrutinib of different concentrations without radiation on Day 9. Scale bar: 100 μm for brightfield pictures. **(C)** Dose-response curves of circularity of HN30 and HN31 organoids under treatment of ibrutinib, acalabrutinib or spebrutinib of different concentrations. Data under treatment of ibrutinib are shown in blue, data under acalabrutinib are shown in blue, and data under spebrutinib are shown in brown. Data of non-irradiated organoids are shown as solid lines, and data of irradiated ones as dashed lines. Data are represented as mean ± SEM. **(D)** Dose-response curves of Day 9 normalized average size of organoids of HN30 under treatment of ibrutinib, acalabrutinib or spebrutinib of different concentrations. Organoid sizes are normalized to those of DMSO control organoids on Day 1. Data under treatment of ibrutinib are shown in blue, data under acalabrutinib are shown in blue, and data under spebrutinib are shown in brown. Data of nonirradiated organoids are shown as solid lines, and data of irradiated ones as dashed lines. Data are represented as mean ± SEM. **(E)** Scatter plots of normalized viability versus Day 9 circularity for HN31 organoids treated with ibrutinib, acalabrutinib or spebrutinib of different concentrations, overlapped with half-translucent Figure 2F. Data are represented as mean ± SEM. Data under treatment of ibrutinib are shown in blue, data under acalabrutinib are shown in blue, and data under spebrutinib are shown in brown. Irradiated data are shown in a slightly different shade compared to nonirradiated ones. Linear regression analysis was done for nonirradiated data of HN31 organoids under treatment of ibrutinib (shown as the green line) or spebrutinib (shown as the brown line). See also **Supplementary Figure 7**.

The effect of different BTK inhibitors on organoid size was also consistent with their potency (**Figure 3B-C, Supplementary Figure 7**), as was the effect on circularity in more invasive HN31 organoids (**Figure 3B, D**). We plotted viability and circularity data for HN31 organoids treated with each of the three BTK inhibitors at various concentrations and observed the same trends: ibrutinib showed significant anti-invasion effects, while spebrutinib had small effects, and acalabrutinib had no effects (**Figure 3E**).

Since acalabrutinib and spebrutinib have superior BTK selectivity compared to Ibrutinib, we conclude that ibrutinib’s effects on HN30 and HN31 organoids are most likely due to its off-target effects.

### Immunofluorescence of HN30 and HN31 Organoids Maps Differences between Ibrutinib, Cetuximab and Ipatasertib

To further investigate how ibrutinib and the EGFR inhibitor cetuximab lead to toxicity in HN30 organoids and inhibit invasion of HN31 organoids, and how the AKT inhibitor ipatasertib promotes invasion of HN30 organoids, we performed immunofluorescence studies of HN30 and HN31 organoids under different treatment conditions (i.e. with or without radiation, treated with 1µM ibrutinib, cetuximab, ipatasertib or DMSO as a negative control). We labeled markers of invasion (F-actin [52], EGFR [41], cytokeratin 14 (K14) [53] and matrix metalloproteinase 2 [MMP2]), epithelial-mesenchymal transition (EMT; E- and N-cadherin, and Vimentin), DNA damage (γH2AX), and proliferation (Ki67), resulting in a detailed view of processes that are affected by the three inhibitors (**Supplementary Figure 8-9**).

In terms of the effects on invasion, we observed that Ibrutinib greatly reduced the formation of F-actin in HN30 organoids but not in HN31, whereas cetuximab and ipatasertib treatments had minimal or no effect, respectively (**Figure 4A, Supplementary Figure 8A**). Additionally, we observed that higher levels of EGFR in untreated HN30 organoids were reduced upon ibrutinib treatment (**Figure 4A**), whereas treatment of HN31 with ibrutinib and cetuximab resulted in increased EGFR staining (**Figure 4A, Supplementary Figure 8A**). These results indicate that inhibitors, especially ibrutinib, have different effects on invasion of HN30 and HN31 organoids, perhaps reflecting distinct origins (primary vs. metastatic).

**Figure 4.**
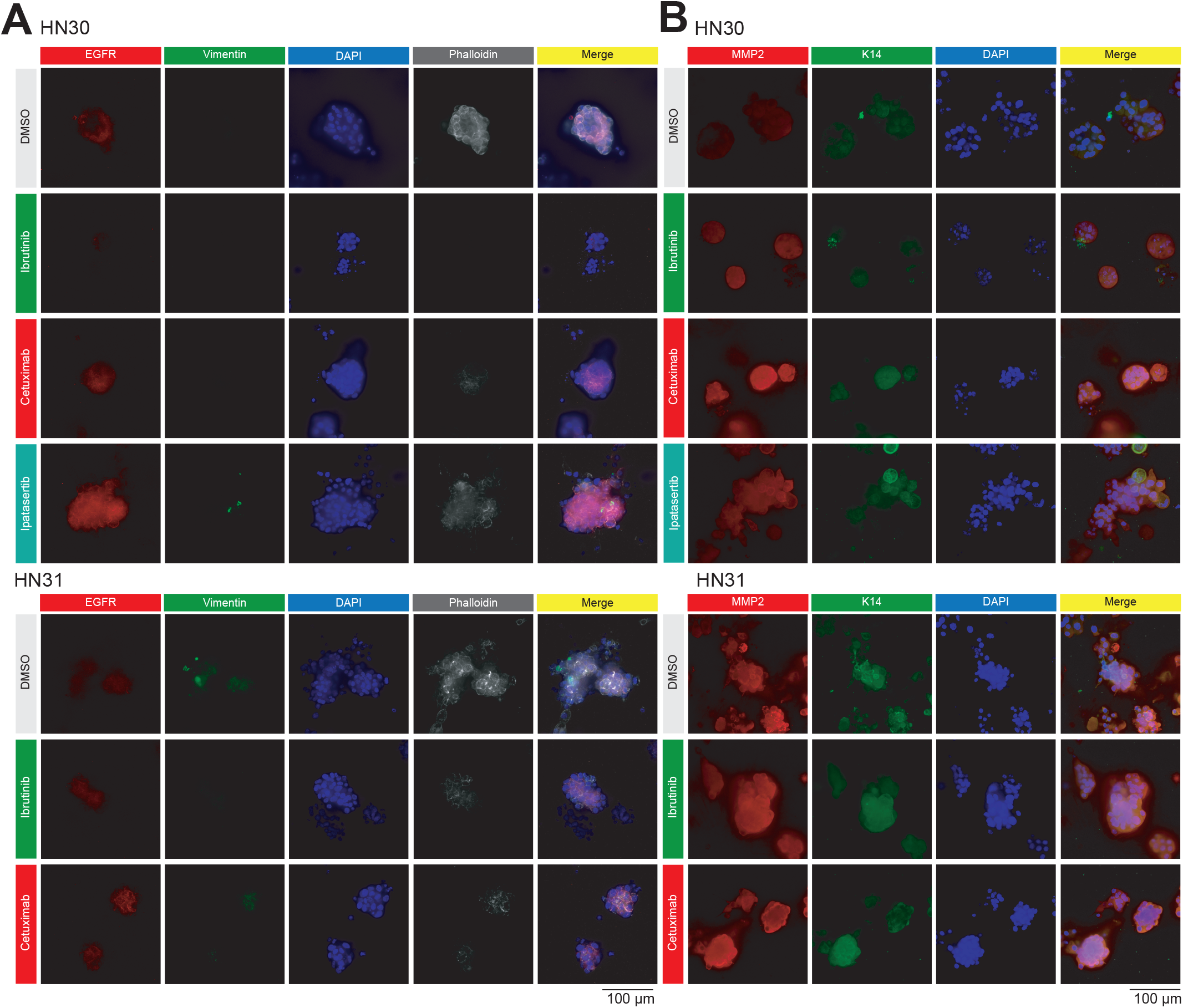
Immunofluorescence Images of HN30 and HN31 Organoids. **(A)** Representative immunofluorescence images showing EGFR (red), vimentin (green), nuclei (DAPI, blue) and F-actin (phalloidin, white) in nonirradiated HN30 organoids treated with DMSO vehicle, ibrutinib, cetuximab or ipatasertib and nonirradiated HN31 organoids treated with DMSO vehicle, ibrutinib or cetuximab. Scale bar: 100 μm for immunofluorescence images. **(B)** Representative immunofluorescence images showing MMP2 (red), K14 (green) and nuclei (DAPI, blue) in nonirradiated HN30 organoids treated with DMSO vehicle, ibrutinib, cetuximab or ipatasertib and nonirradiated HN31 organoids treated with DMSO vehicle, Ibrutinib or Cetuximab. Scale bar: 100 μm for immunofluorescence images. See also **Supplementary Figure 8-10**.

The effects of the inhibitors on EMT were visualized by staining vimentin, a mesenchymal marker, and cadherins. We observed that vimentin only showed expression in few HN30 or HN31 organoids, and showed no specific association with the invading status of cells inside organoids, making it an unreliable marker of invasion in these systems (**Figure 4A, Supplementary Figure 8A**). We also examined E-cadherin. Both HN30 and HN31 organoids showed membrane staining of E-cadherin, and this pattern of staining did not change upon treatment (**Supplementary Figure 9A**). These results suggest that both HN30 and HN31 organoids maintain this epithelial marker, and the three inhibitors we tested (with and without radiation) do not apparently reduce the expression of this marker in HN30 and HN31. Additionally, neither ibrutinib nor cetuximab affected HN30 organoids through DNA damage (as labeled by γH2AX; **Supplementary Figure 10**) or had effects on proliferation (as visualized using Ki67 labeling) in HN30 or HN31 (**Supplementary Figure 9B**). Therefore, EMT, DNA damage and cell proliferation are not substantially perturbed upon treatment with ibrutinib, cetuximab or ipatasertib, and although radiation did increase γH2AX levels indicative of DNA damage (**Supplementary Figure 10B**), treatment with inhibitors did not change these levels.

Overall, immunofluorescence documented the effects of these three targeted therapies on HNSCC organoids, highlighting distinct patterns of response. These results, and the other data presented, provide strong support that our HNSCC organoid-based platform facilitates screening of therapeutic responses, and enables studies of physical and molecular features of organoids that can be used to further refine predictions regarding treatment efficacy.

### Development of HPV+ Patient-Derived Cell Lines and Oropharyngeal Tumor Organoids

To demonstrate that our platform can perform high-throughput analysis of clinical samples, we collected 76 patient samples from 45 patients (mostly male, Caucasian, and 50–70 years old; see **Supplementary Table 2** for a detailed description of our patient cohort). Samples collected included palatine and lingual tonsil (91%), base of tongue (8%), and other subsites (1%). Tumor type was predominantly squamous cell carcinoma (SCC), but also included pleomorphic adenoma, lymphoid hyperplasia, and high-grade dysplasia. Most samples were early-stage HPV+ oropharyngeal squamous cell carcinoma, of which four were used to grow organoids for the high-throughput platform (**Table 1**).

**Table 1.** Details of patients from whom we derived four different organoids.

| CELL/ORGANOID ID | HPV Status | Patient Characteristics<br>(Sex/Age by Decade*/Race) | Site of Dissection |
| --- | --- | --- | --- |
| MS0004RTU | HPV+ (p16+) | Male/60s/White | Right Tonsil |
| MS0006RTU | HPV+ (p16+) | Male/50s/White | Right Tonsil |
| MS0041RTU | Unknown,<br>Reactive follicular<br>hyperplasia<sup>1)</sup> | Male/60s/White | Right Tonsil |
| MS0044RTU | HPV+ (p16+) | Male/50s/White | Right Tonsil |

\* Age by decade of diagnosis via biopsy; 1) p16+ metastases found in a lymph node, while no primary site was identified.

To characterize our chosen patient-derived cell lines (i.e. MS0004RTU, MS0006RTU, MS0041RTU, and MS0044RTU) before forming organoids, we performed the whole-exome analysis of extracted DNA. We observed that each cell line exhibited unique patterns, numbers, and type of mutations. MS0006RTU and MS0041RTU had lower mutational burden (9 and 12 mutations, respectively), than MS0004RTU and MS0044RTU (39 and 104 cases of mutations, respectively) (**Supplemental Table 3**). While mutations in MS-0006RTU were predominantly indels, frameshift or nonframeshift, nonsynonymous single nucleotide variants (SNV) were dominant in MS0004RTU and MS0044RTU. To focus our attention on genes known to be involved in cancer, we compared our results with a list of 447 cancer-relevant genes from the Dana-Farber OncoPanel (POPv3) [54],leading to the identification of 13 cancer-relevant genes with SNVs (**Figure 5A**). For example, we observed a frameshift deletion in ASXL1 (an epigenetic regulator) in MS0041RTU, a nonframeshift deletion in PHOX2B (a transcription factor) in MS0004RTU, and two stop-gain mutations in KMT2D (a lysine methyltransferase) and MDM2 (an E3 ubiquitin ligase) in MS0044RTU. Additionally, MS0006RTU showed a nonsynonymous SNV in GNAS (alpha-subunit of the stimulatory G protein), MS0044RTU had four cases of non-synonymous SNV (i.e. in two kinases (ALK and PIK3CA), a guanine nucleotide exchange factor (ARHGER12), and a histone acetyl transferase (EP300)), and MS0004RTU had two nonsynonymous SNVs, in CYLD (a deubiquitinase) and NR0B1 (a nuclear receptor). The only shared genetic change we detected is a frameshift deletion of the same cytosine in MSH2, a protein involved in DNA mismatch repair, that occurred in both MS-0041RTU and MS0006RTU.

**Figure 5.**
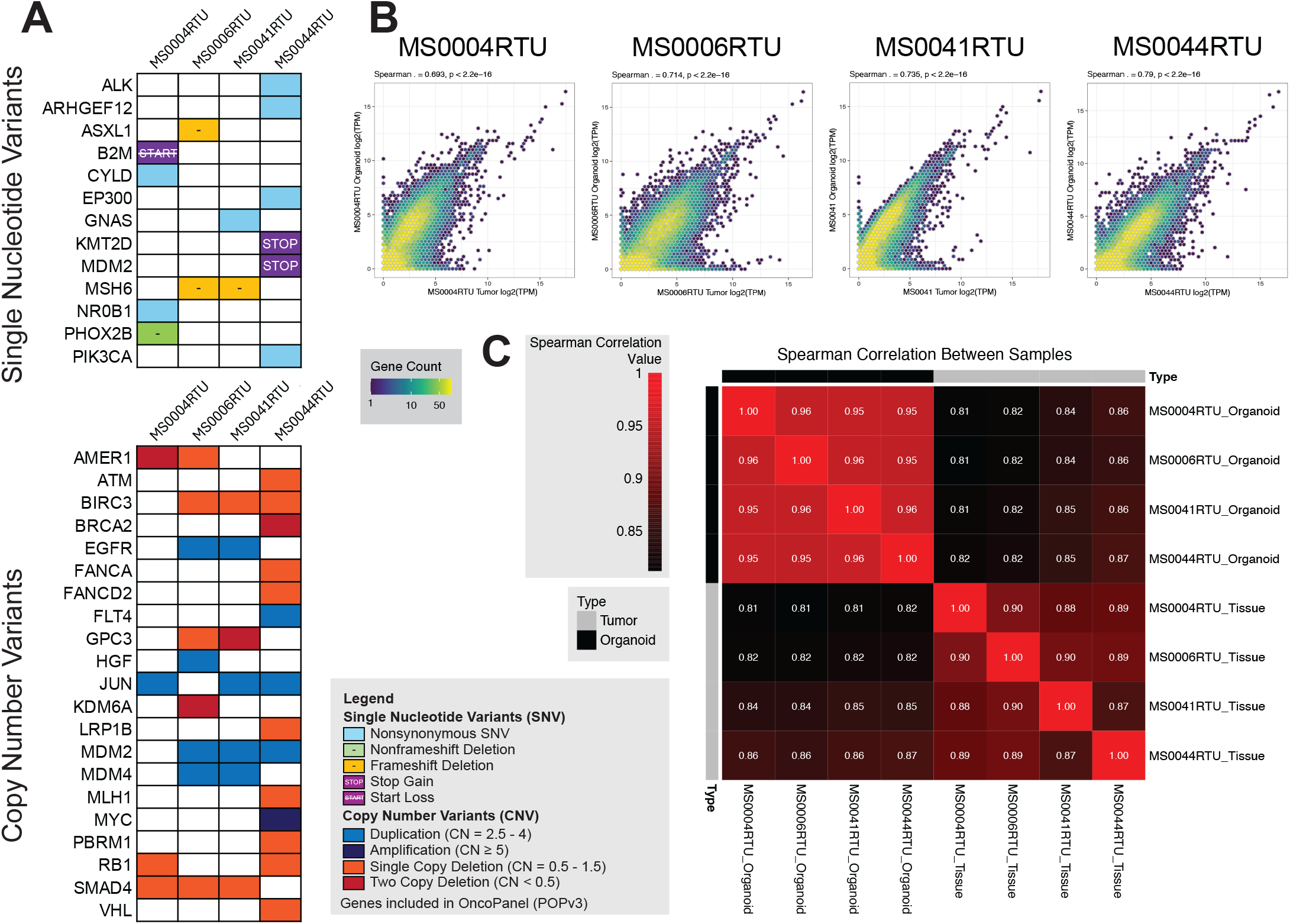
DNA and RNA Sequencing Analysis of Clinical Samples. **(A)** Genomic landscape of clinical samples of MS0004RTU, MS0006RTU, MS0041RTU and MS0044RTU. The table below shows findings from genomic characterization of tumor samples with functional annotation of genetic variants by WANNOVAR and copy number variant calling by DRAGEN. Colors of the heatmap indicate the types of mutations and copy number variants: nonsynonymous single nucleotide variant (SNV), frameshift or nonframeshift deletion, stop gain, start loss, duplication, amplification, single copy deletion and two copy deletion. Genes listed in OncoPanel (POPv3) are shown here. **(B)** Spearman’s rank correlation of RNA abundance comparing RNA transcripts of tumor and organoids derived from MS0004RTU, MS0006RTU, MS0041RTU and MS0044RTU. We found strong correlations between RNA abundance in cells directly from tumor tissue and from organoids grown in vitro. p-values are derived from a two-tailed test of correlation between paired samples. **(C)** Spearman’s rank correlation coefficients comparing RNA transcripts of tumor and organoids derived from MS0004RTU, MS0006RTU, MS0041RTU and MS0044RTU. Genes with expression less than 0.1 transcripts per million are excluded from the analysis.

The four cell lines also exhibited different levels of copy number variants (CNVs) – 484 CNVs in MS-0044RTU, 525 CNVs in MS0006RTU, 642 CNVs in MS0004RTU, and 733 CNVs in MS0041RTU (for full data see **Supplemental Table 3**; for a summary focused on proteins known to be involved in cancer see **Figure 5A**). We observed that several CNVs occurred in multiple clinical samples, such as copy number deletions in BIRC3 (an inhibitor of apoptosis), SMAD4 (a tumor suppressor), GPC3 (a heparan sulfate proteoglycan), AMER1 (Wnt pathway regulator), and RB1 (a tumor suppressor); and copy number duplications in JUN (a transcription factor), MDM2, EGFR and MDM4 (an MDM2 homolog). Of note, MS0044RTU exhibited high level of oncogenic MYC (CN = 6.2) amplification, suggesting that this tumor could be most difficult to treat. Overall, our DNA sequencing analysis shows that patient-derived cell lines exhibit clear genetic differences, making them suitable for validating our platform. Therefore, we used these cell lines to generate four patient-derived organoids, which showed to recapitulate gene expression in cancer tissue as measure by transcriptomics (**Figure 5B-C**). These validations indicate that our patient-derived organoids are valuable models for testing drug responses under different treatment conditions.

### Drug Screening of Patient-Derived HPV+ Oro-pharyngeal Tumor Organoids

To examine the effect of radiation on patient-derived organoids, we exposed them to a radiation gradient (0 to 8 Gy) and observed that all four organoids were sensitive to radiation (less than 50% viability at 4 Gy for all organoids (**Figure 6A**). Since their sensitivities to radiation between 2 or 4 Gy were various, we used 1 Gy for MS-0044RTU and MS0004RTU and 2 Gy for MS0041 and MS0006RTU in the following drug screening. While irradiated organoids of other clinical samples had ideal viability for measurement of radiosensitizing abilities of therapy agents (MS0004RTU at 87.9%, MS0044RTU at 94.7%, and MS0006RTU at 87.3%) at their designated doses, the viability of irradiated MS0041RTU organoids was as low as 64.2% at 2 Gy (**Figure 6B**). Therefore, we chose to exclude MS0041RTU from our screens for radiosensitizers and focused on using MS0004RTU, MS0044RTU, and MS0006RTU under their respective radiation doses for those experiments.

**Figure 6.**
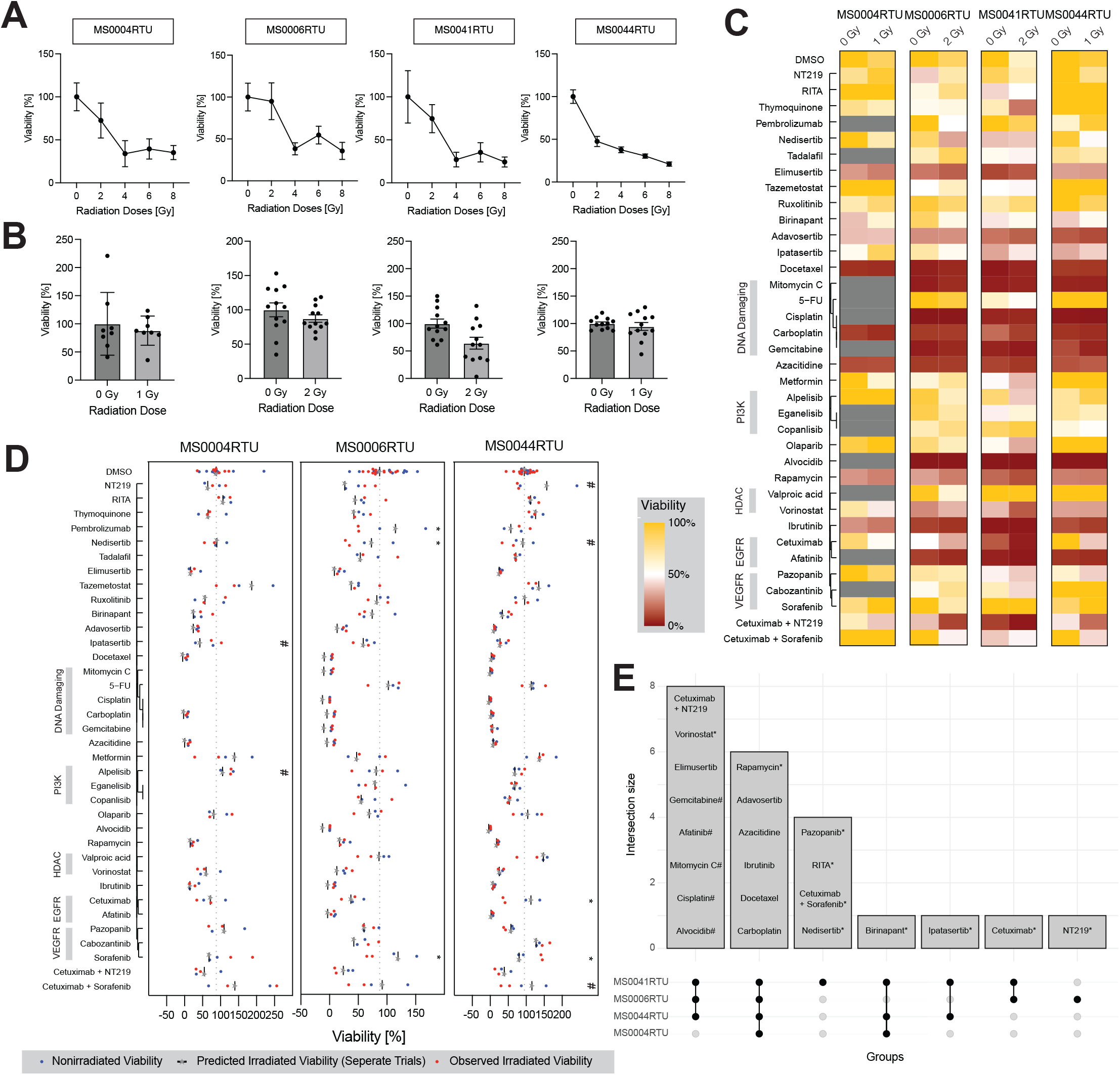
Clinical Samples Viability and Combination Effects with Radiation. **(A)** Dose-response curves of organoid viability of MS-0004RTU, MS0006RTU, MS0041RTU and MS0044RTU exposed to radiation of different intensities in the radiation gradient assay. Data are represented as mean ± SEM. **(B)** Normalized viability % of DMSO control organoids of MS0004RTU, MS0006RTU, MS0041RTU and MS0044RTU with and without radiation in high-throughput drug screening. Data are represented as mean ± SEM. **(C)** Heatmap of high-throughput screening results of MS0004RTU, MS0006RTU, MS0041RTU and MS0044RTU for 33 therapy agents and 2 combinations with Cetuximab, with or without radiation of specific doses for each cell line. The color represents the normalized cell viability %. The drugs are clustered using the Jaccard distance based on common protein targets. **(D)** Scatter plots of high-throughput screening results for organoids of MS0004RTU, MS0006RTU, MS0041RTU and MS0044RTU assessing the combination effects of therapeutic agents with radiation. Blue dots represent the normalized cell viability % of individual wells without radiation. Grey stars and black bars represent predicted post-radiation viability, calculated using a linear interaction model based on the agent-specific effect, derived from the average viability of all nonirradiated wells within the same trial. Red dots represent observed viability % of wells treated with both the therapeutic agent and radiation. Synergistic effects are shown when red dots lie to the left of the black bar, which indicates lower than expected viability. Student’s t-test was used to compare predicted and observed irradiated values. *: q < 0.01. For MS0004RTU, #: p < 0.1; for MS0044R-TU, #: 0.01 ≤ q < 0.1. **(E)** Upset plot showing therapy agents effective against organoids of MS0004RTU, MS0006RTU, MS0041RTU and MS0044RTU. Criteria: leading to normalized viability < 50% and q-value < 0.1. #: This therapy agent is not screened in MS0004RTU. *: The therapy agent is not effective in SCC154 according to the aforementioned criteria. See also **Supplementary Figure 11**.

Before studying the effects of drugs on radiation sensitivity, we first wanted to establish the baseline and screen MS0006RTU, MS0041RTU, and MS0044RTU organoids with the same compound library as the one used above. For MS-0004RTU organoids, where we had a limited number of available cells, we selected 22 drugs and two cetuximab combinations for drug screening by removing therapy agents inhibiting overlapping targets and/or pathways, and those found to be toxic. We found that the following drugs were effective against all clinical samples screened, leading to viability lower than 50% and q-value smaller than 0.1: (1) chemotherapy agents cisplatin, carboplatin, gemcitabine, mitomycin and docetaxel; (2) pan-CDK inhibitor alvocidib; (3) EGFR inhibitor afatinib; (4) BTK inhibitor ibrutinib; (5) DNA methylation inhibitor azacitidine; (6) Wee1 inhibitor adavosertib; and (7) mTOR inhibitor rapamycin (Figure 6C). Additionally, a subset of compounds was effective against 3 of 4 samples (anti-ATR inhibitor elimusertib (not significantly in MS0004RTU); anti-HDAC vorinostat (not in MS0004RTU); anti-IAP inhibitor birinapant (not in MS0006RTU); and the combination of cetuximab and NT219 (not in MS-0004RTU)), with AKT inhibitor ipatasertib effective against MS0041RTU (viability 31.2%) and MS-0044RTU (viability 32.3%), and nearly so against MS0004RTU (viability 54.6%) (**Figure 6C**). In general, our patient-derived organoids displayed a similar drug sensitivity profile to the HPV+ cell line SCC154, with rapamycin, vorinostat and ipatasertib being exceptions (active on patient-derived organoids only). MS0041RTU was sensitive to several drugs in the library (18 out of 33 drugs), including the DNA-PK inhibitor nedisertib, the HDM2 inhibitor and TP53 activator RITA, the VEGFR and PDGFR inhibitor pazopanib, and the combination of cetuximab and sorafenib – none of which was effective in the other three clinical samples. We also note that MS0006RTU was specifically sensitive to IRS1/2 and STAT3 inhibitor NT219.

To identify compounds that act as radio-sensitizers, we compared predicted viability and observed viability of organoids exposed to their respective radiation doses. This analysis showed that cetuximab is a powerful radiosensitizer in MS-0044RTU, with radiation transforming this originally resistant case into one that is susceptible to cetuximab (**Figure 6D**). All cetuximab combinations were also very effective when further supplemented with 1 Gy radiation. In MS0006RTU, the PD-1 inhibitor pembrolizumab (observed viability 50.3% compared to predicted viability 114.7%, q = 0.000683) and the multikinase inhibitor sorafenib (observed viability 119.3% compared to predicted viability 70.0%, q = 0.0041) were significant radiosensitizer, with sorafenib and cetuximab combination having milder effects (observed viability 46.8% compared to predicted viability 91.8%, p = 0.10, q = 0.167) (**Figure 6D**). Finally, metformin (observed viability 61.2% compared to predicted viability 138.8%, p = 0.21) sensitized MS0004RTU to radiation treatment (**Figure 6D**), whereas nedisertib radiosensitized both MS0006RTU and MS0044RTU (**Figure 6D**). Additionally, in MS-0044RTU, combination of cetuximab and NT219 or sorafenib in MS0044RTU was found to be synergistic (normalized viability 44.4%, β6 estimate 0.2563, p = 0.0156), while the combination of radiation, cetuximab and NT219 displayed antagonism (β7 estimate 1.339, p = 0.0115) (**Supplementary Figure 11**). We also observed that radiation nullifies the overgrowth under treatment with NT219, resulting in a viability of 77.1% from the predicted 157.2% (**Figure 6D**).

Taken together, our patient-derived organoids allowed us to map viability under different treatment conditions. As shown, this type of test could rapidly yield insights into which drugs (or drug combinations) are likely to be effective in each patient, and whether complementing drug treatment with radiation therapy is likely to be more effective (we discuss these points in more detail below).

### Image Analysis of Patient-Derived Organoids

As the final step in our platform performance assessment and validation, we conducted image analysis of two patient-derived organoids (MS0006RTU and MS0044RTU). In both cases, we observed that drug treatment as well as drug + radiation treatment result in small changes in organoid size (**Figure 7A-D, Supplementary Figure 12**). Additionally, these organoids are generally spheroidic and do not exhibit an invasive phenotype (as visualized by the appearance of protrusions and decrease in circularity), including upon treatment (**Figure 7A-B, E-F, Supplementary Figure 12**). In MS0044RTU, cetuximab led to a size decrease in the absence of radiation, with the normalized average observed organoid size of 1.508 in comparison to the predicted 2.438 (according to linear regression; q = 0.37) (**Figure 7D, Supplementary Figure 13**). In MS0006RTU, sorafenib led to a decrease in size, with a normalized average organoid size of 2.253, compared to the predicted value of 2.905 (q = 0.72, and a large variation across wells) (**Figure 7C, Supplementary Figure 13**). In all cases, radiation diminished these effects (**Figure 7C, Supplementary Figure 13**). Collectively, these data support that combining our platform with brightfield imaging opens opportunities for a more complete analysis of how drugs and/or radiation affect the size and shape of organoids, including patient-derived organoids as shown here.

**Figure 7.**
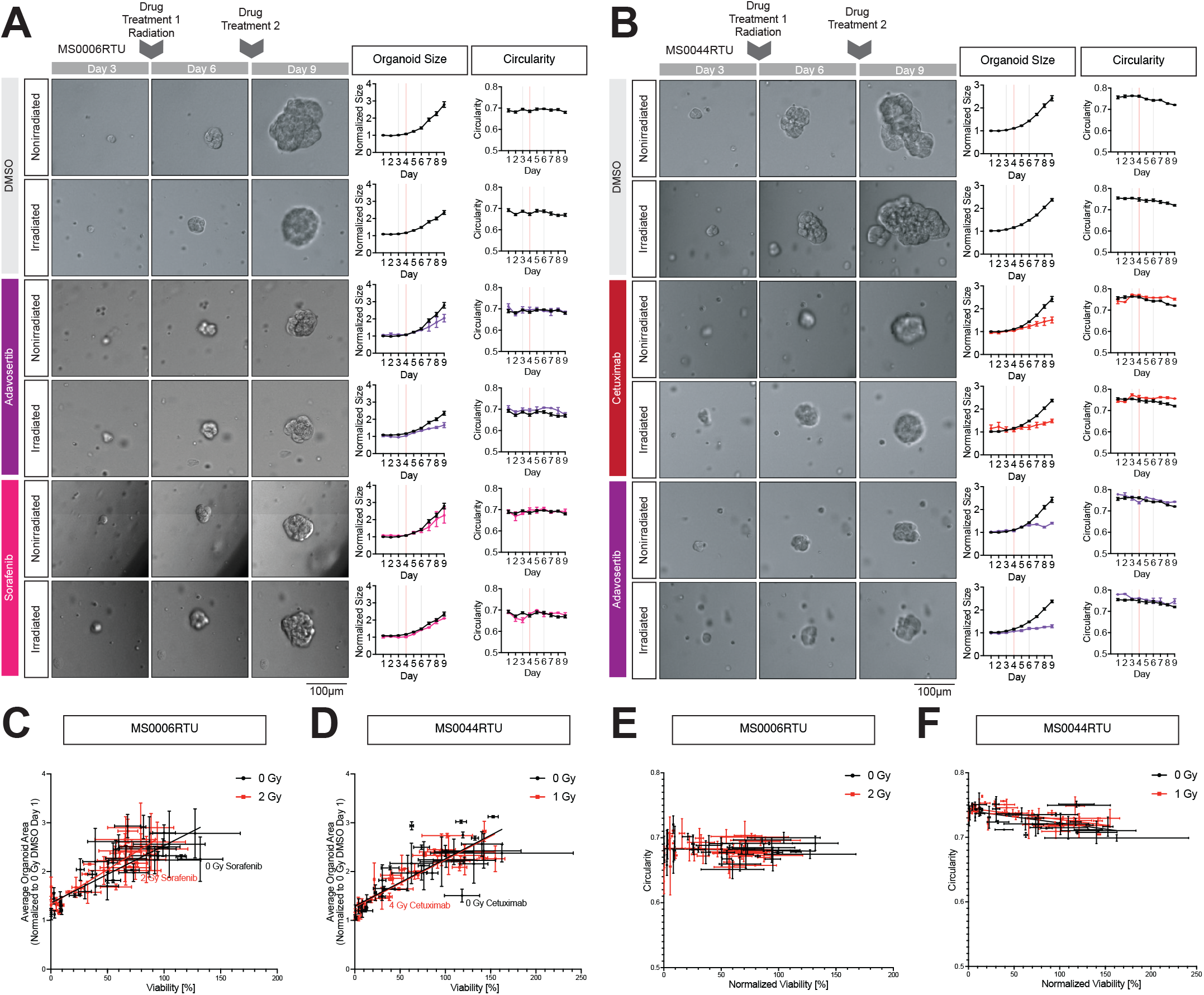
Image Analysis Results of Patient-Derived MS0006RTU and MS0044RTU Organoids. **(A-B)** Representative brightfield images of organoids of MS0006RTU **(A)** and MS0044RTU **(B)** in culture on Day 3 (Column 1), Day 6 (Column 2) and Day 9 (Column 3). Organoids were treated with therapy agents on Day 3 and Day 6, and radiation on Day 4, after imaging. Growth (Column 4) and changes of morphology of organoids (Column 5) was tracked over time by segmenting in-focus organoids in the brightfield images using a machine learning–based pipeline. Growth was measured with the normalized organoid size every day, which is calculated by normalizing average cross-sectional area of all organoids across all wells of the same treatment of the same cell line across all trials to that of DMSO without radiation measured on Day 1 of culture. Data of control organoids are denoted with black lines. Data of treated organoids are denoted with respective colors. Data are represented as mean ± SEM. Morphology was measured with average circularity of all organoids across all wells of the same treatment of the same cell line across all trials. Scale bars: 100 μm for brightfield pictures. **(C-D)** Scatter plots of normalized viability % versus Day 9 normalized organoid size for MS0006RTU **(C)** and MS0044RTU **(D)**. Each dot denotes one therapy agent or combination. Data without radiation are denoted with black and data with radiation are denoted with red. Data are represented as mean ± SEM. Linear regression analysis were done for nonirradiated and irradiated data separately, with the black line the linear regression line of nonirradiated data and the red line that of irradiated data. **(E-F)** Scatter plots of normalized viability versus Day 9 circularity for MS0006RTU **(E)** and MS0044RTU **(F)**. Data are represented as mean ± SEM. Linear regression analysis was done for nonirradiated and irradiated data separately, with the black line the linear regression line of nonirradiated data and the red line that of irradiated data. See also **Supplementary Figures 12 – 13**.

## Discussion

HNCs are a group of cancers with heterogeneous mechanisms and management strategies due to their diverse etiologies [3,9,55]. The current standard of care, i.e. surgery followed by radiation therapy, carries risks associated with the toxicity of high-dose radiation. In more advanced disease setting, especially unresectable cases of the HPV+ subtypes, standard of care also includes chemotherapy (e.g. cisplatin), further increasing risks of toxicity and adverse effects [56]. Therefore, using personalized treatment regimens with optimized doses and less toxic alternatives may overcome these challenges and offer more effective, safer and tolerable treatments for HNCs [20,57,58]. However, as mentioned, HNCs are heterogenous, and finding the best match between patient-specific HNC and therapy remains difficult. In this study, we built an automated high-throughput platform that uses patient-derived organoids to identify therapeutics that are most likely to exhibit efficacy in patients. We showed that our platform can find candidates that are effective in killing malignant cells, as well as those that cooperate with radiotherapy to reduce the minimum functional dose of radiation, and those that are effective in preventing cancer invasion and metastasis to significantly improve patient prognosis [16,17].

Using our platform we demonstrated that different organoids exhibit distinct patterns of drug sensitivity (e.g. HPV+ SCC154 organoids are more sensitive to radiation than HPV-HN30 and HN31, in agreement with previous studies) [59], which is driven by differences in their genetics. For example, HN30 organoids are more sensitive to birinapant, a SMAC mimetic and IAP inhibitior, than HN31, which agrees with a previously reported observation that HN31 cells are resistant to SMAC mimetics [60]. Furthermore, HN31 are more sensitive to Docetaxel than HN30, which is consistent with previous findings that microtubule inhibitors are more effective against morphologically more invasive “sarcomatoid” or “mixed sarcomatoid-epithelial” cancer cases such as HN31 [24]. Similarly, we noticed that Wee-1 inhibitor adavosertib is more effective against HPV+ cases than HPV-ones, consistent with previous studies [61,62].

However, there are also several results that differ from the existing literature. Previous studies showed that ATR inhibitor elimusertib effects were independent HPV+ or HPV-status [63,64]. In contrast, we find that elimusertib eliminates SCC154 and most of HPV+ patient-derived organoids, but not HPV-organoids (HN30 and HN31). This discrepancy might be due to HN30 and HN31-specific resistance to elimusertib that is not present in other HPV-subtypes. In another example, our HPV+ MS0044RTU and MS0004RTU organoids are resistant to MDM2 inhibitor RITA, which does not match results from a previous study showing that RITA can suppress HPV+ tumors [65]. Although some of these observations will need future follow up to dissect factors that impacted these differences, the results we present demonstrate that our organoid-based platform can identify key differences between HPV− or HPV+ tumors and their sensitivity to different drugs in an automated and high-throughput manner. This suggests that our platform is highly compatible with an unmet medical need, i.e. matching patients with an appropriate personalized therapy.

This is further supported by our results using patient-derived organoids generated from HPV+ oropharyngeal tumor samples of four patients. We showed that patient-derived organoids shared sensitivity to multiple drugs we screened, many of which were also effective against HPV+ SCC154 organoids (e.g. alvocidib, afatinib, ibrutinib, azacitidine, adavosertib, elimusertib, and several chemotherapy agents). On the other hand, we also noticed that HPV+ organoids are resistant to several generally toxic agents, such as mTOR inhibitor rapamycin, HDAC inhibitor vorinostat, SMAC mimetic birinapant, and AKT inhibitor ipatasertib. We propose that HPV infection is directly linked to the observed resistance. For example, HPV viral oncoproteins E6 and E7 affect PI3K/AKT/mTOR signaling pathway [66,67], potentially causing resistance to inhibitors of this pathway, Rapamycin and Ipatasertib [68–70]. Additionally, HPV E7 was proposed to lead to resistance against Rapamycin by activating Phospolipase D and degrading Rb [71], indicating that multiple, HPV-dependent mechanisms may be in play. Furthermore, HPV E6 and E7 also regulate histone modifications of infected cells [72], which could potentially interfere with vorinostat. However, previous in vitro studies showed that vorinostat reduces growth of HPV+ cervical cancer cell lines by reducing expression of E6 and E7 [73–75], and HPV+ HNSCC cell lines by inhibiting p63 isoform ΔNp63α, which is upregulated with HPV infection [76,77]. Therefore, although reconciling these results will require additional studies, we propose that the difference may be caused by higher concentrations of Vornistat used in earlier work [76]. We were somewhat surprised to note that Birinapant is effective in HPV+ patient-derived organoids given that IAP signaling pathways are mostly normal in HPV+ HNSCC cancer cell lines [78]. These results highlight that, being more physiologically relevant, patient-derived organoids may detect drug sensitivities that 2D cell-based assays fail to detect. Overall, we present a comprehensive dataset that integrates drug sensitivity profiles with specific patient-derived organoids, thus providing the foundation for future studies of how differences in sensitivities correlate to differences in genomic landscapes of these clinical samples.

The same platform can be leveraged to identify agents with radiosensitizing activity. We found radiosensitizing activity for the multi-kinase inhibitor pazopanib in SCC154, the FDA-approved EGFR inhibitor cetuximab in MS0044RTU, the multi-kinase inhibitor sorafenib selectively in MS0006RTU, and the DNA-PK inhibitor nedisertib in both MS0044RTU and MS0006RTU. These findings are consistent with prior preclinical and clinical reports for cetuximab [67,79], sorafenib [80,81], and nedisertib [82–84]. Pazopanib has previously been reported to radiosensitize sarcoma xenograft models through endothelial dysfunction [85], although its direct effects on tumor cells and its clinical relevance as a radiosensitizer remain unclear [86,87]. Importantly, the activity of pazopanib, cetuximab, sorafenib, and nedisertib differed across the four patient-derived organoid models analyzed, suggesting substantial inter-patient variability in radiosensitivity responses. In this context, our platform may provide a scalable approach to identify patient-specific radiosensitizer candidates through functional screening.

Remarkably, we also show how combining of our platform with imaging and ML-based image analysis can be used to go beyond viability as the end-point readout and discover agents that inhibit invasion and metastasis by measuring and comparing morphology of organoids. The drug screening of HN31 organoids, derived from an invasive (metastatic) cell line, showed that EGFR inhibitors afatinib and cetuximab, pan-PI3K inhibitor copanlisib and BTK inhibitor ibrutinib change the morphology of those organoids from sprouting (invasive) into spheroidal (non-invasive). These results are in agreement with what is known about the roles of EGFR, PI3K and BTK in metastasis. For example, EGFR mutations play a role in brain metastasis of non-small-cell lung cancer, and EGFR inhibitors significantly improve overall survival in these patients [88,89]. In HNSCC, overexpression of EGFR is associated with poor prognosis, and treatment with cetuximab in addition to platinum-based therapies prolonged survival of recurrent and metastatic patients [43]. Similarly, PI3K is involved in actin cytoskeleton reorganization via both downstream AKT and Rac signaling pathways [44]. Previous trials showed that copanlisib treatment led to partial response in metastatic breast cancer cases [45], again supporting the conclusions that emerged from our platform about afatinib, cetuximab, copanlisib and ibrutinib as therapies that could inhibit cancer invasion and improve outcomes for metastatic HNSCC patients. Importantly, our screening platform also detected therapies that promote invasion, such as AKT inhibitor ipatasertib, and to lesser extent rapamycin, inhibitor of mTOR which is downstream of AKT. This suggests that the use of AKT inhibitors could be detrimental to HNSCC patients, urges caution, and provides another example of how our platform can assist in clinical decision making.

In addition to the use cases described above that illustrate how our platform can advance personalized medicine and facilitate matching patients with individualized therapies, our results also suggest that the platform is an effective approach for drug repurposing work. Among 34 drugs screened in our study, only 6 have been approved for HNC therapy, while 16 have been approved for therapy of other cancers, and 4 that are approved for other diseases (**Table 1**). One such example of a repurposing candidate is Ibrutinib, a BTK inhibitor approved for treating blood cancers [50,51]. We observed that ibrutinib effectively inhibits the growth of organoids of HN30, SCC154 and all tested clinical samples, radiosensitizes HN30 and HN31, and inhibits the invasion of HN31 organoids. Importantly, various preclinical trials show that ibrutinib is also effective in killing solid tumors, including HNC [46], further supporting our conclusions that Ibrutinib repurposing for treating HNSCC should be further considered. Of note, when compared to more BTK-selective second-generation inhibitors (acalabrutinib and spebrutinib) ibrutinib had distinct effects, suggesting that BTK inhibition is not significantly involved in ibrutinib’s toxic, radiosensitizing, or invasion-inhibiting effects on HNC. Outside of BTK, ibrutinib also targets EGFR, HER2, HER4, BLK, BMX, JAK3, and other molecules [90], and further studies are needed to pinpoint which of these proteins is the most relevant target for ibrutinib in the context of HNSCC.

### Limits of the Current Platform

The linear interaction model we use in the study to connect effects of radiotherapy and different anti-cancer drugs works best when neither of the two treatments is too toxic. Therefore, for highly toxic drugs, our approach is not able to show synergetic effect with radiotherapy [91,92]. One example of this issue is Ibrutinib, which is not shown to be a radiosensitizer in HN30 because it is already toxic at the concentration of 1 µM (**Figure 1B-C**). In this case, lowering the concentration eventually revealed radiosensitizing activity of Ibrutinib (**Figure 3A**); however, this issue may also explain why our study does not show radiosensitizing effects of highly toxic chemotherapy agents like cisplatin. Therefore, for potential radiosensitizing candidates, further assays with different concentrations and radiation doses are recommended to measure combination effects with radiation.

Also important is the determination of a radiation dose for the drug screening assays, since an inadequately selected radiation dose cannot be used to measure synergism between radiation and various therapeutic agents. We recommend using a radiation dose that leads to a normalized viability of over 90%.

We also acknowledge that using circularity of individual organoids as a measure of invasiveness, which works well for collective invasion [93], may not work for other invasion modes like mesenchymal-like or amoeboid-like individual dissemination. In that case, other methods of measurement should be used.

## Conclusion

Here, we developed the first automated, high-throughput patient-derived HNSCC organoid platform for performing drug sensitivity screening. The platform is compatible with both viability assays and imaging-based analysis and supports use of radiotherapy together with all other major classes of drug treatments (i.e. chemotherapy, targeted therapy, and immunotherapy). With this organoid-based high-throughput platform, we have the ability to establish a personalized therapeutic pipeline for patients with advanced HNSCC, maximizing responses to radiotherapy and targeted agents to improve outcomes, avoid modulators that promote tumor invasion, and find new therapies through drug repurposing efforts.

## Acknowledgments

We thank the Technology Center for Genomics & Bioinformatics (TCGB) for assistance with sequencing. We thank the Department of Medicine Statistics Core of UCLA Health for advice on statistical analysis.

## Methods

### Cell lines

HN30 and HN31 cell lines are provided by Dr. Jeffrey N. Myers of MD Anderson Cancer Center, while the SCC154 cell line is from ATCC. HN30, HN31 and SCC154 cell lines were cultured in Dulbecco’s modified Eagle’s medium with high glucose, L-glutamine, phenol red and sodium pyruvate (DMEM, Gibco™, 11995-065) supplemented with 10% heat-inactivated fetal bovine serum (FBS, Gibco 16140-071 and A56708-01) and 1% antibiotic-antimycotic (Gibco 15240-062), optionally with 0.2% MycoZap™ Prophylactic (Lonza, VZA-2032). Only cells passaged within 10 generations were used in the experiments. All three cell lines were authenticated by short tandem repeat profiling using the GenePrint 10 kit (Laragen, TransnetYX) with leftover cells after experiments or at the end of the 10th passage. In order to distinguish HN30 and HN31, their DNA was extracted using the PureLink™ Genomic DNA Mini Kit (Invitrogen™, K182002) according to the manufacturer’s protocols. After, the sequence surrounding Exon 5–6 of TP53, where mutations specific to HN31 (A161S, C176F) are located, were amplified with PCR using primers (forward: cgctagt-gggttgcagga, reverse: cactgacaaccacccttaac [55], produced by Integrated DNA Technologies), DreamTaq PCR Master Mixes (2X) (Thermo Scientific™, K1071), and a MiniAmp™ Plus Thermal Cycler (Applied Biosystems™, A37835) following the manufacturers’ protocols. The following PCR programs were used: 1 cycle of initial denaturation (95°C, 1 min), 35 cycles of denaturation (95°C, 30 s), annealing (65°C, 30 s), extension (72°C, 1 min), and 1 cycle of final expension (72°C, 5–15 min). After verification with gel electrophoresis with UltraPure™ Agarose (Invitrogen™, 16500-100), UltraPure™ TAE Buffer, 10X (Invitrogen™, 15558-026), SYBR™ Green I Nucleic Acid Gel Stain (Invitrogen™, S7563), and BlueJuice™ Gel Loading Buffer (Invitrogen™, 10816015), PCR products were then purified and sequenced with Sanger sequencing by Laragen and TransetYX. Mycoplasma was detected with real time PCR with Mycoplasma-specific primers and probes by Laragen and TransnetYX. For some experiments, preliminary mycoplasma detection was performed after cell passage and before the experiments with MycoAlert® PLUS Mycoplasma Detection Kit (Lonza, LT07-710) according to the manufacturer’s protocols.

### Clinical sample collection and processing

Tumor tissues were collected from consented adults undergoing transoral robotic surgery and processed (IRB approved protocols #11-002858, #15-001395 and #19-000947). Tissue samples were harvested by the Translational Procurement Core Laboratory under the direction of Dr. Maie St. John and Dr. Alice Soragni. The protocol for collecting and processing tumor tissue has been previously described [28,29,31,32]. Clinical samples were minced manually with a scalpel and digested with collagenase IV (200 U/mL) to dissociate into individual cells or clusters, from which red blood cell were lysed with Ammonium Chloride Solution (Stem Cell Technology, 07850). Cells were then strained through a 100 μm filter. Counting and viability assessment were implemented with a Cellometer Auto 2000 (Nexcelom).

### Whole exome sequencing and analysis

Whole-exome sequencing analysis was performed by the Technology Center for Genomics & Bioinformatics (TCGB) of the University of California Los Angeles (UCLA). Genomic DNA was extracted with DNeasy Blood & Tissue Kit (Qiagen). The KAPA HyperPrep Kit (Roche) was used to construct an Illumina-compatible library. KAPA HyperCap v3.5 for Human WES (Roche) was used for KAPA Target Enrichment with Probes. The libraries were sequenced using Novaseq X Plus PE 150bp. Data quality checks were performed on Illumina SAV. Demultiplexing was performed using Illumina Bcl2fastq v2.19.1.403 software. For data analysis, reads were aligned to Human HG38 (Homo sapiens [1000 Genomes] hg38 v5) genome using Illumina DRAGEN Somatic version 4.4 software [95]. DRAGON Somatic version 4.4 Tumor vs Normal mode was used for secondary analysis. WANNOVAR was used for functional annotation of genetic variants found in hard-filtered. vcf generated by DRAGEN Somatic [96]. We filtered hits with a list of 447 cancer relevant genes tested by OncoPanel (POPv3) [54].

### RNA sequencing and analysis

RNA sequencing results between cells collected from clinical samples and from organoids were compared. For organoids used for RNA sequencing, 100,000 cells were seeded in 70 μL hydrogel with a 3:4 mixture of organoid culture media and Matrigel (Corning, 354234) around perimeter of each well of 24-well plates. After 30-minute incubation at 37°C, 1 mL organoid culture media was added to each well. Organoids were cultured for 2 weeks, and media were added on Day 3, 6, 9 and 11. On Day 13, organoids were collected after release with dispase. RNA sequencing and analysis were performed by the TCGB of UCLA. Total RNA was extracted from samples using the RNeasy Mini kit (Qiagen). Libraries for RNA-Seq were prepared with the KAPA Stranded RNA-Seq Kit with the RiboErase Kit. The workflow consists of depletion of rRNA via hybridization with complementary DNA oligonucleotides, followed by treatment with RNase H and DNase and RNA fragmentation. First strand cDNA synthesis using random priming followed by second strand synthesis converting cDNA:RNA hybrid to double-stranded cDNA (dscDNA), and incorporates dUTP into the second cDNA strand. cDNA generation is followed by A-tailing, adaptor ligation and PCR amplification. Different adapters were used to multiplex samples in a single lane. Sequencing was performed on an Illumina NovaSeq X Plus using a PE 2×150bp run. Data quality checks were performed on Illumina SAV. Demultiplexing was performed with Illumina Bcl2fastq v2.19.1.403 software.

### 3D printing plasma masks

The design and 3D printing of custom well masks for plasma treatment of glass-bottom plates (Cellvis, P96-1.5H-N) were conducted as previously described [30]. Briefly, Inventor 2020 (Autodesk) was used to design masks for the ring area around the edge of each well for treatment. The design, saved as a.STL file, was imported into the PreForm (FormLabs) software to arrange the parts and command the Form3B 3D printer (FormLabs) to complete printing with the Biomed Amber resin (FormLabs, RS-CFG-BMAM-01). After printing, parts were cleaned with compressed gas dusters (Office Depot, 110284), washed twice in isopropanol, air-dried for at least 30 minutes, and cured for an additional 30 minutes at 70 °C in the Form Cure UV oven (FormLabs, FH-CU-01).

### Bioprinted 3D organoids

High throughput drug screening for organoids was conducted as previously described [30]. In short, organoids were bioprinted with a commercially available temperature-controlled extrusion bioprinter (Matribot, Corning, 6150) onto a glass-bottom plate. A python script was made to convert a Microsoft Office Excel worksheet with information of bioprinted patterns (rings), including size (5.5mm in diameter) and volume (10µL), to be bioprinted (30 wells for radiation gradient assay, 60 wells for clinical sample drug screening or BTK inhibition assay, or 96 wells for cell line drug screening of the whole 96 well plates). Full G-code files were generated using the coordinates for each well to command the bio-printer to achieve the desired single-layer geometry. Succinctly, on Day 0, after cell passage (for cell lines) or tissue dissociation (for clinical samples), a single-cell suspension with a concentration of 300 cells per µL for cell lines and 500 or 1,000 cells per ul for clinical samples, was placed in a 3:4 mixture of organoid culture media and basement membrane–based hydrogel (Cultrex™ Basement Membrane Extract, Pathclear for cell lines, R&D systems, 3432-005-01; Matrigel for clinical samples, Corning, 354234) prepared and kept on ice. After brief vortexing, the mixture was transferred into a 3 mL syringe, which, after the attachment of a 18G needle (CELLINK, NZ5180505001), was inserted into the bioprinter where the printhead was maintained at a print temperature of 2°C. Concurrently, 3D-printed plasma masks were inserted into the 96 well plate and pressed in contact with the glass surface using rubber bands. The plate was subsequently treated in a plasma cleaner (PE-25, Plasma Etch) for 90 seconds. After removal of the masks, the glass-bottom plate was placed on the platform of the bioprinter.The printer was calibrated by moving the needle to the bottom right corner of the plate, and the X and Y coordinates were recorded before Z calibration. Post-calibration, the bioprinter was primed by extruding the gel through the needle until a droplet formed. Each ring-structure in a 96-well plate was printed in approximately five minutes. After printing, constructs were incubated at 37°C for at least 30 minutes to solidify the matrix, followed by the addition of pre-warmed 100 μL of organoid culture media using an automated fluid handler (Microlab NIMBUS, Hamilton or epMotion 96, Eppendorf). Organoid culture media was prepared with Pneumacult™-Ex (STEM-CELL, 05008) supplemented with 0.1% Hydrocortisone Stock Solution (STEMCELL, 07925). For cell lines, 1% antibiotic-antimycotic was added with 0.2% MycoZap Prophylactic added optionally. For clinical samples, 1% antibiotic-antimycotic or 0.2% Primocin (Invivogen, ant-pm-2) was added. Additionally, each multiwell plate was imaged daily using a high-content microscope (Celigo, Nexcelom) every day.

### Media change and drug screening

On Day 3 and Day 6, media change or drug treatments were implemented with the therapy agents mentioned previously. Old media is removed and new media, laden with therapy agents or vehicles mentioned below, is added using an automated fluid handler. The positive control is 10 μM Staurosporine and the negative control is 1% DMSO. Each plate contains its own positive and negative controls for normalization. For the radiation gradient assay, where positive control and other wells continued to be cultured without DMSO, the positive control was optional. Drugs and DMSO were dispensed into pre-warmed organoid culture media with a non-contact dispenser (Flexdrop™ iQ, Perkin-Elmer). However, cetuximab, pembrolizumab, and valproic acid were added manually. For drug screening of cell lines or clinical samples, therapy agents listed in **Supplementary Table 1**, other than cisplatin (50 μM), carboplatin (50 μM) and thymoquinone (10 μM), were managed at 1 μM. For the BTK inhibition assay, ibrutinib, acalabrutinib and spebrutinib were managed at concentrations of 0.2, 0.5, 1, 2, 5, 10, 20 and 50 μM. For organoids used for immunostaining, 1μM Ibrutinib, cetuximab and ipatasertib were treated manually.

### Irradiation

On Day 4, organoids were irradiated at RT with an experimental X-ray irradiator (Gulmay Medical Inc). The X-ray beam was operated at 300 kV and 10 mA and hardened using 4 mm Be, 3 mm Al, and 1.5 mm Cu filters and calibrated using NIST-traceable dosimetry. For the preliminary radiation gradient assay, 2 or 3 plates were bioprinted for each cell line or clinical sample. Each plate was divided into three regions: control, low radiation (2 / 6 / 12 Gy) and high radiation (4 / 8 / 16 Gy). First, a metal sheet was used to cover only the control region to absorb radiation, and the plate was irradiated until the low and high radiation regions were exposed to 2 / 6 / 12 Gy radiation. Then, the metal sheet covered both the control and low radiation regions, and the high radiation region was exposed to an additional 2 / 2 / 4 Gy of radiation. For drug screening and BTK inhibition assays, each plate was divided into two halves: control and irradiated regions. During radiation treatments, plates with control regions were covered with the metal sheet until the irradiated region was exposed to a dose optimized according to the results of the radiation gradient assay.

### ATP assay

On Day 9, organoid viability was assessed using an ATP assay with an automated fluid handler, as previously described [28,29]. A PBS wash was completed thereafter, and the organoids were released from the hydrogel with dispase. After five minutes of vigorous shaking at 80 rpm, 30 μL (for cell lines) or 75 μL (for clinical samples) CellTiter-Glo reagent (Promega, G968B) was added to each well. After a 30-minute RT incubation, followed by five minutes of vigorous shaking at both ends, luminescence was measured using a SpectraMax iD3 plate reader (Molecular Devices).

### Immunofluorescence and confocal imaging

For immunofluorescence of HN30 and HN31 organoids at the end of culture, gels were fixed in 10% formalin with 1% Glutaraldehyde (50 wt. %, Sigma-Aldrich), permeabilized with DPBS with 0.5% Triton X-100, and blocked in 1x PBS with 10% heat-inactivated fetal bovine serum and 0.2% Triton X-100. Primary antibodies were incubated overnight at 4°C at the dilutions listed below in antibody diluent prepared with PBS with 2% heat-inactivated fetal bovine serum and 0.2% Triton X-100. Secondary antibodies coupled to Alexa Fluor 488, 562 (Invitrogen) were incubated for 2 hrs at room temperature, along with Phalloidin (Alexa Flour Plus 647, Invitrogen, 1:400 dilution) and Hoechst 33342 (Invitrogen, 1:1000 dilution). Primary antibodies were E-cadherin (Invitrogen, HECD-1, 2 µg/mL), N-cadherin (Invitrogen, PA529570, 1:200), EGFR (Invitrogen, H11, 1:100), Vimentin (Invitrogen, PA5-27231, 1:200), γH2AX (R&D Systems, MAB3406, 1:200), phosphor-EGFR (Tyr1068, Invitrogen, 44-788G, 1:100), MMP2 (Invitrogen, 436000, 2 µg/mL), K14 (Invitrogen, PA5-28002, 1:100), BTK (Santa Cruz Biotechnologies, 7F12H4, 1:100), and Ki67 (Abcam, SP6, 1:200). Confocal images were acquired using a ECHO Revolution confocal microscope with a 40x Olympus oil objective. For z-stacks, 1 μm spacing was used. Red-green-blue (RGB) images were assembled using Fiji software and custom scripts.

### Quantification and statistical analysis

Statistical analysis was carried out with GraphPad Prism 10. The Department of Medicine Statistics Core of UCLA Health also provided advice on statistical analysis.

### Quantification of viability

Cell viability values are ATP luminescence assay values normalized to the nonirradiated group treated with vehicle (DMSO) and expressed as a percentage (%). We used R and ggplot2 package to plot heatmaps. Drugs were clustered by their overlapping targets with the Jaccard distance function in R. To show the effects of therapy agents in each case, multiple unpaired t-tests with Welch’s correction were used to compare cell viability values between the control and treated groups for each therapy agent. The q-value was calculated with p-values adjusted to a 1.00% false discovery rates (FDR) calculated using a two-stage set-up (Benjamini, Krieger, and Yekutieli). Hits were determined by the criterium that the q-value < 0.005. Effects of therapy agents between HN30 and HN31 were also compared similarly, and t-tests were used to compare cell viability values across treated groups for each therapy agent in the two cell lines. To select therapy agents that are more effective against SCC154 than against HN30 or HN31 cell lines, a two-way ANOVA test was performed on cell viability values from treated groups of each therapy agent across the three cell lines, with SCC154 values set as the control. The p-values were adjusted with Dunnett statistical hypothesis testing, and the alpha threshold was set at 0.05 (95% confidence interval). The hits were determined with the criteria that (1) the cell viability of SCC154 is smaller than both of the values of HN30 and HN31, and (2) the adjusted p-values between SCC154 and HN30 and HN31 are both smaller than 0.0001.

### Quantification of combination effects

The Linear Interaction Effect model was used to analyze combination effects between cetuximab and NT219 or sorafenib, and between radiation and therapy agents [58]. Briefly speaking, the effect of each therapy is calculated as follows:

*Viability (Therapy)=1-Effect (Therapy)*

It is assumed that if there were no interaction effects between two therapies, i.e. they were independent of each other, the predicted effect would be *Predicted Effect (Therapy A,Therapy B)=Effect (Therapy A)+Effect (Therapy B)*. Then, we can get the predicted viability of the combination:

*Predicted Viability (Therapy A,Therapy B) =1-Predicted Effect (Therapy A,Therapy B) =1-Effect (Therapy A)-Effect (Therapy B) =1-(1-Viability (Therapy A))-(1-Viability (Therapy B)) =Viability (Therapy A)+Viability (Therapy B)-1*

Then, the combination effects of two therapies can be determined by comparing the predicted viability and observed viability which is measured from ATP luminescence assay values of organoids under two therapies: if observed viability is smaller than predicted viability, then the two therapies are found to be synergetic; if observed viability is larger than predicted viability, then the two therapies are found to be antagonist. The significance of combination effects was analyzed by doing Student’s t-tests between normalized viabilities of each well under two treatments and predicted viabilities of each trial (for cell lines), or, for clinical samples, the predicted viability of the trial with predicted standard deviation calculated with *Predicted σ (Therapy A,Therapy B) = √(σ*^*2*^*(Therapy A) + σ*^*2*^ *(Therapy B))*.

To analyze combination effects of cetuximab and radiation with NT219 or sorafenib, multiple linear regression was performed with GraphPad Prism 10 following its guidelines (https://www.graphpad.com/guides/prism/latest/curve-fitting/reg_how-to-multiple-regression.htm). Briefly speaking, organoid viability values with or without radiation, cetuximab, or NT219 or sorafenib were analyzed with multiple linear regression, with a formula of *Viability= β0 * Intercept+ β1 * Radiation + β2 * Cetuximab + β3 * NT219 or Sorafenib+ β4 * Radiation : Cetuximab + β5 * Radiation : NT219 or Sorafenib + β6 * Cetuximab : NT219 or Sorafenib+ β7 * Radiation : Cetuximab : NT219 or Sorafenib*. Here, parameters β4, β5, and β6 denotes the combination effects of two therapies between colons, and β7 denotes the combination effect of radiation, cetuximab and NT219 or sorafenib together. Those values are negative when two or three therapies concerned have synergetic effects, and they are positive when the therapies have antagonist effects. For each parameter, a p-value was calculated after a Student’s t-test of the comparison between a model with this parameter and a model without this parameter. For BTK inhibition assays, Student’s t-tests were implemented between normalized viabilities of nonirradiated and irradiated groups, and p-values were calculated.

### Quantification of image analysis

Starting on Day 1, we captured brightfield images of organoids daily using a high-content microscope (Celigo, Nexcelom), scanning three focal planes per well and generating whole-well images in TIF format at a resolution of 1 μm/pixel. The images were then segmented and quantified with a previously reported machine learning–based technology [31,32]. In summary, we trained a U-Net convolutional neural network model [97] derived from a model pretrained on the ImageNet dataset [94,98]. We trained over 80 epochs using a cross-entropy loss function, with a training dataset of 50 images of HN31 organoids and a size of 512 x 512 μm featuring diverse organoid morphologies of 17 sites of 2 or 3 focal planes. We segmented organoids in the training dataset with Segment Anything for Microscopy (µSAM) [99]. The generated masks were modified manually in GIMP to include invading sprouts which were frequently omitted by µSAM. To avoid measuring the same organoid across focal planes, we compared images of the three focal planes and only marked the most in-focus organoids. The model generated masks of segmented organoids, marked in black on a white background, enabling quantification of organoid size and morphology using particle analysis in Fiji [100]. A macro script was written to perform particle analysis in batch. Sizes of organoids were quantified as projected areas of particles. Morphologies were quantified as circularity: *Circularity= (4π * Area)/Perimeter*^*2*^

Circularity is inversely related to invasiveness. A value of 1.0 indicates a perfect circle, while a value near 0 indicates a polygon with a lot of long sprouts. For each well, both the average area and circularity of an individual organoid were calculated by averaging the values of all organoids across three focal planes. Then for each condition, means and standard deviations were calculated from all wells across all trials of the case. Organoid growth was calculated by normalizing the average individual organoid area with the value of nonirradiated DMSO wells on Day 1.

### Linear Regression Analysis

To decouple the effects of therapy agents on viability from their effects on organoid size or circularity, a simple linear regression analysis was performed for each case using GraphPad Prism 10. Normalized viabilities and organoid sizes or circularities of each condition were plotted on a scatter plot. Linear regression functions were calculated separately for irradiated and nonirradiated groups. The Chow method of F-test was applied to each dataset to determine whether the slopes of regression lines were significantly different from zero, following the Graph-Pad Prism guidelines (https://www.graphpad.com/guides/prism/latest/curve-fitting/reg_comparingslopesandintercepts.htm). A p-value smaller than 0.05 implies a significant deviation from zero. The slopes and intercepts were compared between nonirradiated and irradiated groups, and groups of interest, to see whether the regression lines of those groups were significantly different from each other. The comparison was performed using an F-test, which evaluates whether the two datasets share a common slope. If the p-value for slope comparison was smaller than 0.05, it was implied that the two regression lines had significantly different slopes. If the slopes were not significantly different (p ≥ 0.05), an additional test for differences in intercepts was performed to determine whether the lines were vertically shifted relative to each other. If the 2nd p-value for slope comparison was greater than 0.05, it was implied that the two regression lines were not significantly different from each other. If the 2nd p-value for slope comparison was smaller than 0.05, it was implied that the two regression lines were parallel to each other [101]. This analysis is equivalent to an Analysis of Covariance (ANCOVA).To identify therapy agents that affect the size or circularity of organoids independent of toxicity, predicted size or circularity for each drug was calculated across all trials using the normalized viability of the well and a linear regression function without radiation. Then, multiple unpaired t-tests with Welch correction were implemented to compare measured and predicted size or circularity of each therapy agent. Hits were determined if the p-values were smaller than 0.01. To determine the influence of radiation treatment onto effects of therapy agents on organoid size or circularity, the distance between measured and predicted size or circularity of each therapy agent was compared between the nonirradiated and irradiated conditions.

