## Supplementary Material for "A Bioprinted Head and Neck Cancer Organoid-Based Platform for Evaluating Multimodal Therapies"

### Supplementary Figure

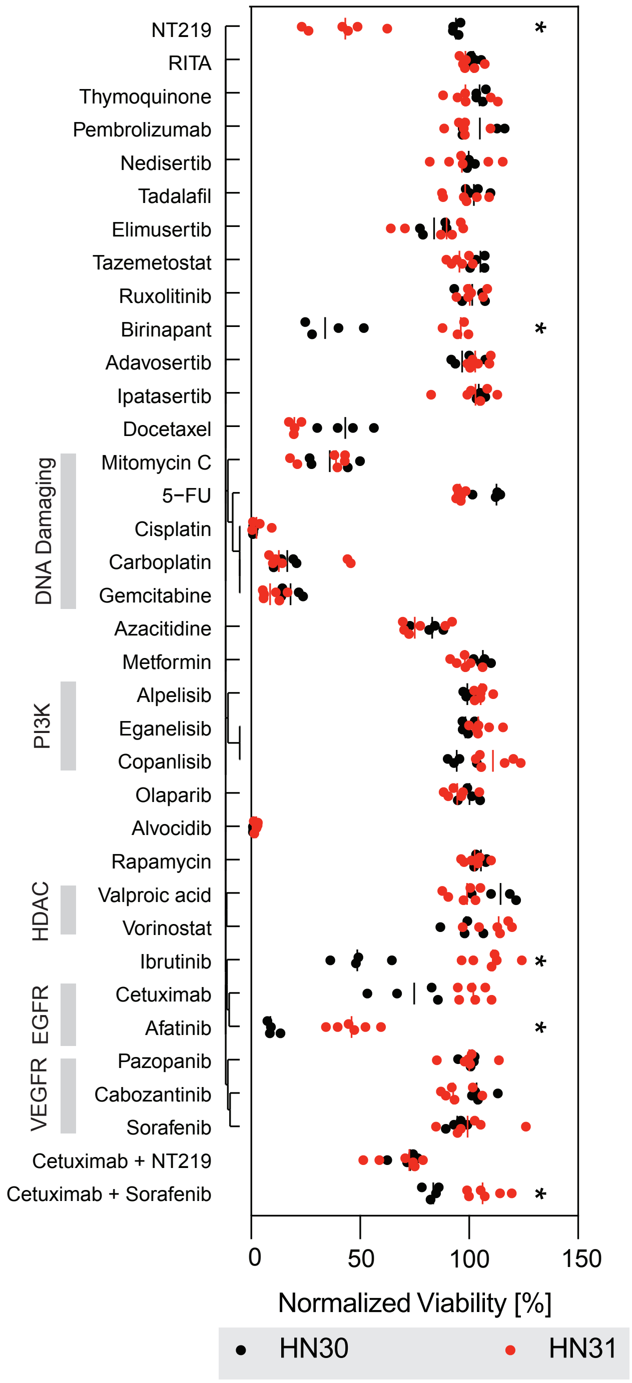


**Supplementary Figure 1.** Related to **Figure 1. Comparison of Viabilities of HN30 and HN31 Organoids Without Radiation.** Scatter plot of high-throughput screening results of HN30 and HN31 without radiation. Each dot represents the normalized cell viability % of one well. The bars represent average normalized cell viability % of all wells of the same treatment of the same cell line. HN30 results are labeled black and HN31 are labeled red. *: $q < 0.01$

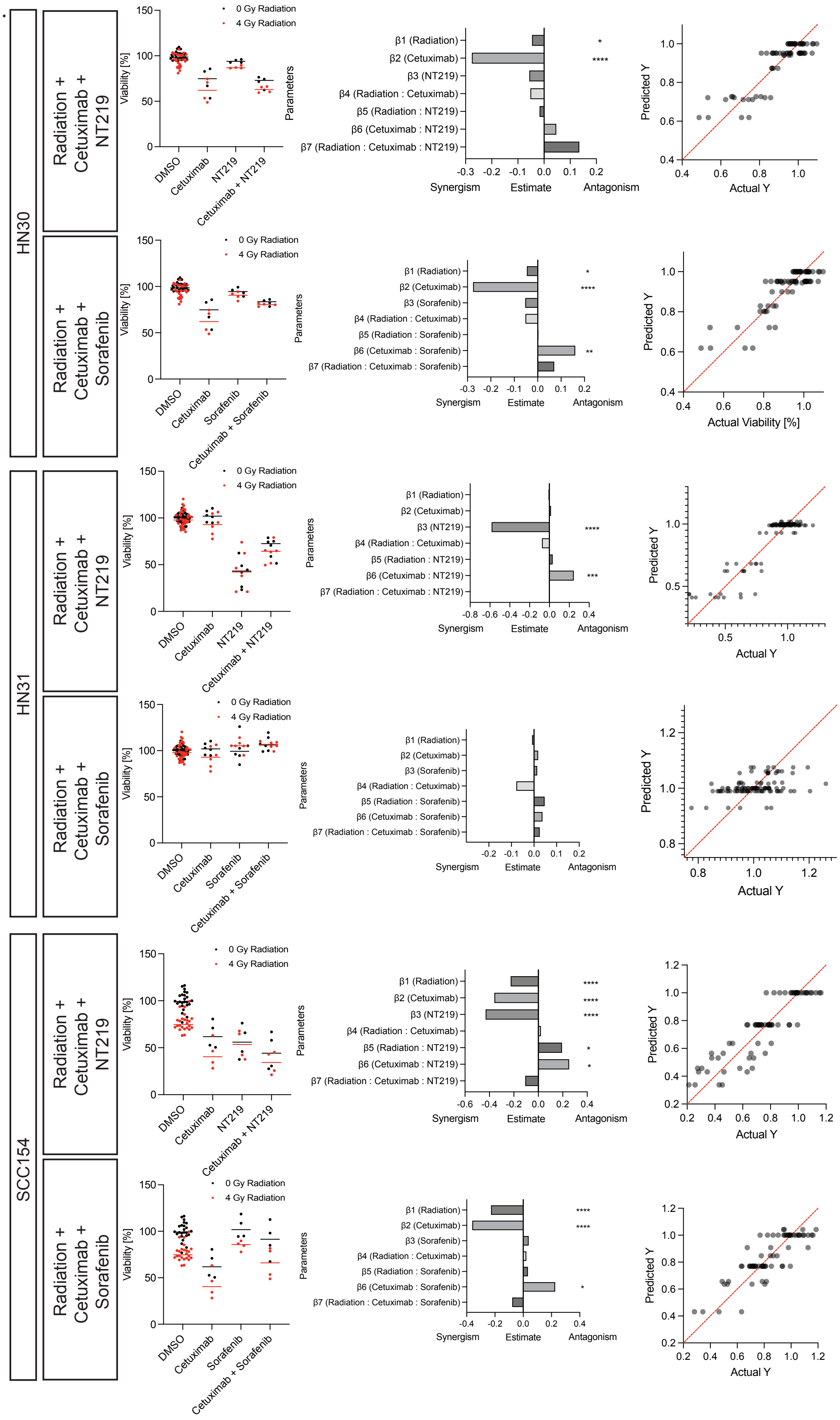
HN30
HN31
SCC154
Radiation + Cetuximab + NT219
Radiation + Cetuximab + Sorafenib
0 Gy Radiation
4 Gy Radiation
Viability [%]
DMSO
Cetuximab
NT219
Cetuximab + NT219
Sorafenib
Cetuximab + Sorafenib
Parameters
β1 (Radiation)
β2 (Cetuximab)
β3 (NT219)
β4 (Radiation : Cetuximab)
β5 (Radiation : NT219)
β6 (Cetuximab : NT219)
β7 (Radiation : Cetuximab : NT219)
β3 (Sorafenib)
β5 (Radiation : Sorafenib)
β6 (Cetuximab : Sorafenib)
β7 (Radiation : Cetuximab : Sorafenib)
Synergism
Estimate
Antagonism
Predicted Y
Actual Y
Actual Viability [%]

**Supplementary Figure 2.** Related to **Figure 1. Combination effects of Cetuximab and NT219 or Sorafenib in cell lines. Left columns:** Normalized viability % of organoids of HN30, HN31 and SCC154 treated with DMSO vehicles, Cetuximab, NT219 or Sorafinib, or combination with Cetuximab and NT219 or Sorafenib, with or without radiation. Data not under radiation are shown in black, and data under radiation are shown in red. **Middle columns:** Data of organoid viability of HN30, HN31, or SCC154 under various treatments were analyzed with multiple linear regression, with a formula of Viability = β0 * Intercept + β1 * Radiation + β2 * Cetuximab + β3 * NT219 + β4 * Radiation : Cetuximab + β5 * Radiation : Agent + β6 * Cetuximab : Agent + β7 * Radiation : Cetuximab : Agent. Colons denote combination effects among two or three treatments. Agent is NT219 or Sorafenib. Bar plots of β1–6. *: 0.0021 ≤ p-value < 0.0332, **: 0.0002 ≤ p < 0.0021:, ***: 0.0001 ≤ p < 0.0002:, ****: p < 0.0001. **Right columns:** The scatter plot of the actual Y values (measured viability, X axis) and predicted Y values (predicted viability, Y axis). The more the data scatter along the line of identity (the red line), the more useful the model is.

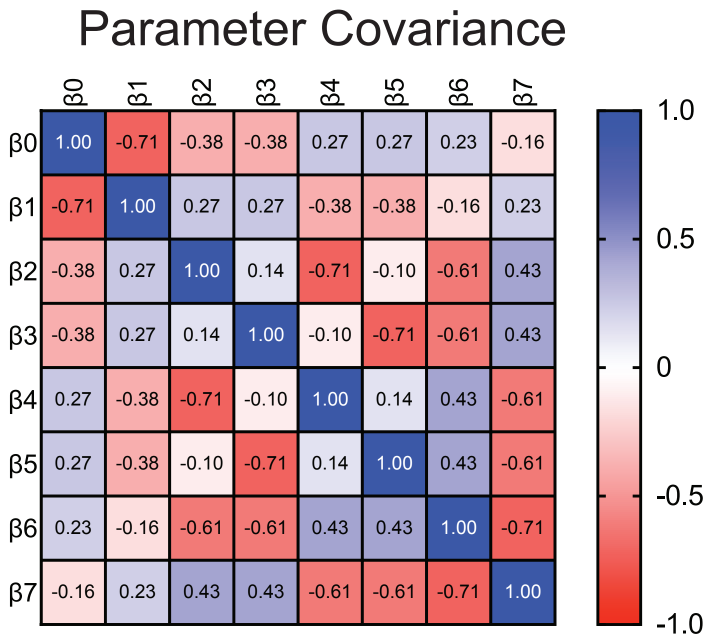


**Supplementary Figure 3.** Related to **Figure 1. Parameter covariance matrix in multiple linear regression used in Supplementary Figure 2.**

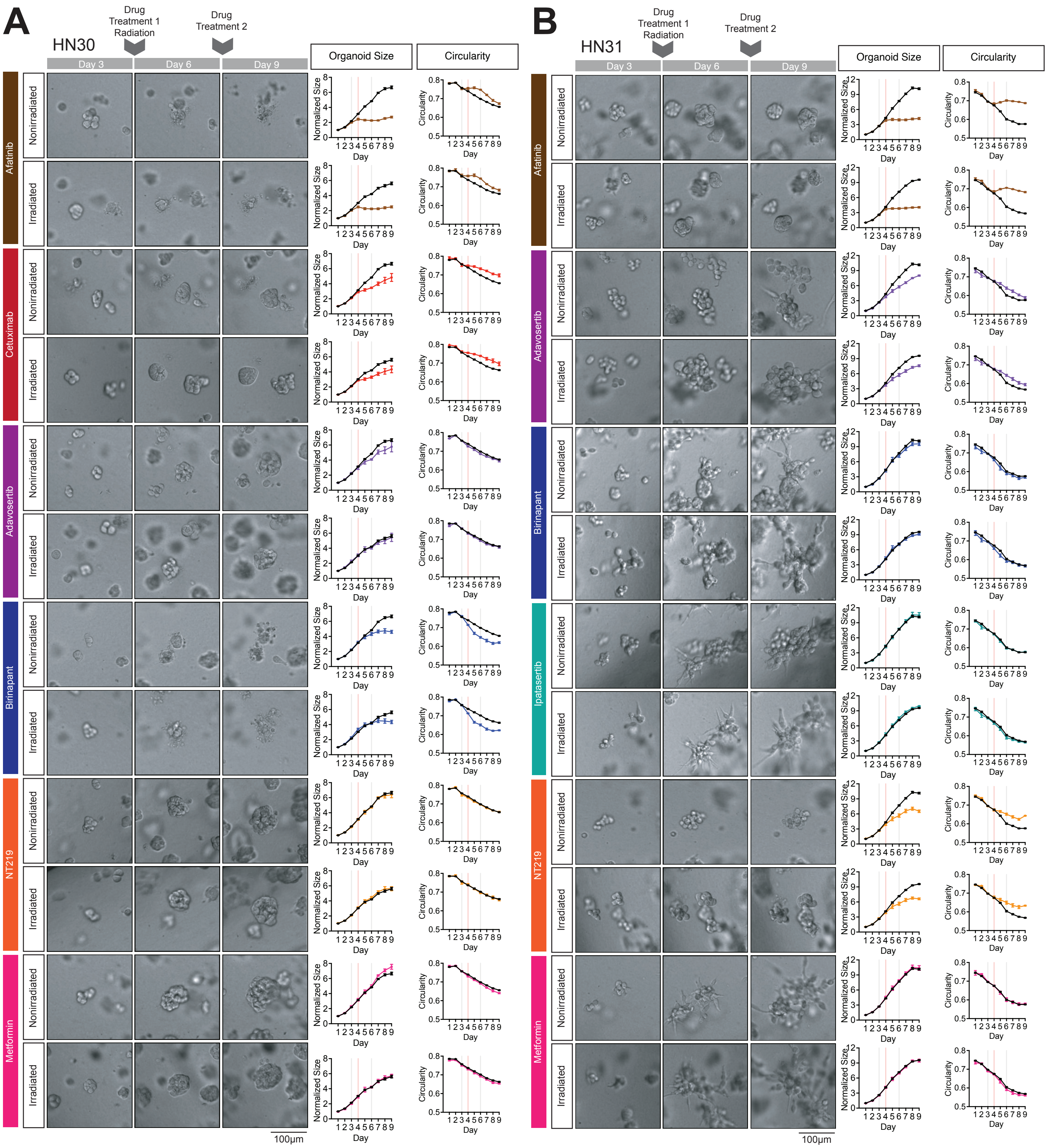


**Supplementary Figure 4.** Related to **Figure 2. Images of organoids and line charts of HN30 and HN31.**

**(A-B)** Supplementary representative brightfield images of organoids of HN30 **(A)** and HN31 **(B)** in culture on Day 3 (Column 1), Day 6 (Column 2) and Day 9 (Column 3). Organoids were treated with therapy agents on Day 3 and Day 6, and radiation on Day 4, after imaging. Growth (Column 4) and changes of morphology of organoids (Column 5) was tracked over time by segmenting in-focus organoids in the brightfield images using a machine learning–based pipeline. Growth was measured with the normalized organoid size every day, which is calculated by normalizing average cross-sectional area of all organoids across all wells of the same treatment of the same cell line across all trials to that of DMSO without radiation measured on Day 1 of culture. Data of control organoids are denoted with black lines. Data of treated organoids are denoted with respective colors. Data are represented as mean ± SEM. Morphology was measured with average circularity of all organoids across all wells of the same treatment of the same cell line across all trials. Scale bars: 100 μm for brightfield pictures.

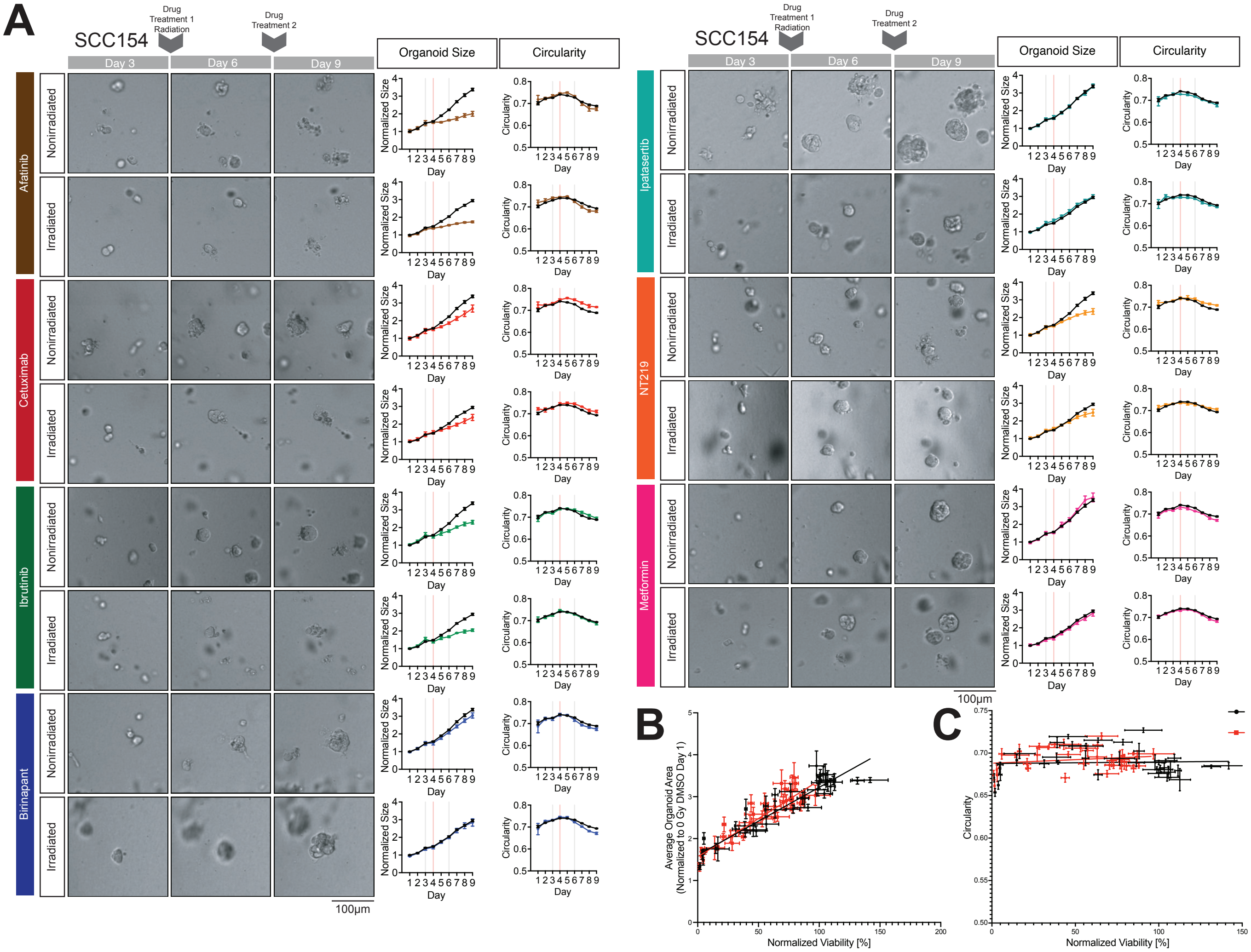


**Supplementary Figure 5.** Related to **Figure 2. Images of organoids, line charts and image analysis results of SCC154. (A)** Supplementary representative brightfield images of organoids of SCC154 in culture on Day 3 (Column 1), Day 6 (Column 2) and Day 9 (Column 3). Organoids were treated with therapy agents on Day 3 and Day 6, and radiation on Day 4, after imaging. Growth (Column 4) and changes of morphology of organoids (Column 5) was tracked over time by segmenting in-focus organoids in the brightfield images using a machine learning–based pipeline. Growth was measured with the normalized organoid size every day, which is calculated by normalizing average cross-sectional area of all organoids across all wells of the same treatment of the same cell line across all trials to that of DMSO without radiation measured on Day 1 of culture. Data of control organoids are denoted with black lines. Data of treated organoids are denoted with respective colors. Data are represented as mean ± SEM. Morphology was measured with average circularity of all organoids across all wells of the same treatment of the same cell line across all trials. Scale bars: 100 µm for brightfield pictures. **(B)** Scatter plots of normalized viability % versus Day 9 normalized organoid size SCC154. Each dot denotes one therapy agent or combination. Data without radiation are denoted with black and data with radiation are denoted with red. Data are represented as mean ± SEM. Linear regression analysis were done for nonirradiated and irradiated data separately, with the black line the linear regression line of nonirradiated data and the red line that of irradiated data. **(C)** Scatter plots of normalized viability versus Day 9 circularity for SCC154. Data are represented as mean ± SEM. Linear regression analysis was done for nonirradiated and irradiated data separately, with the black line the linear regression line of nonirradiated data and the red line that of irradiated data.

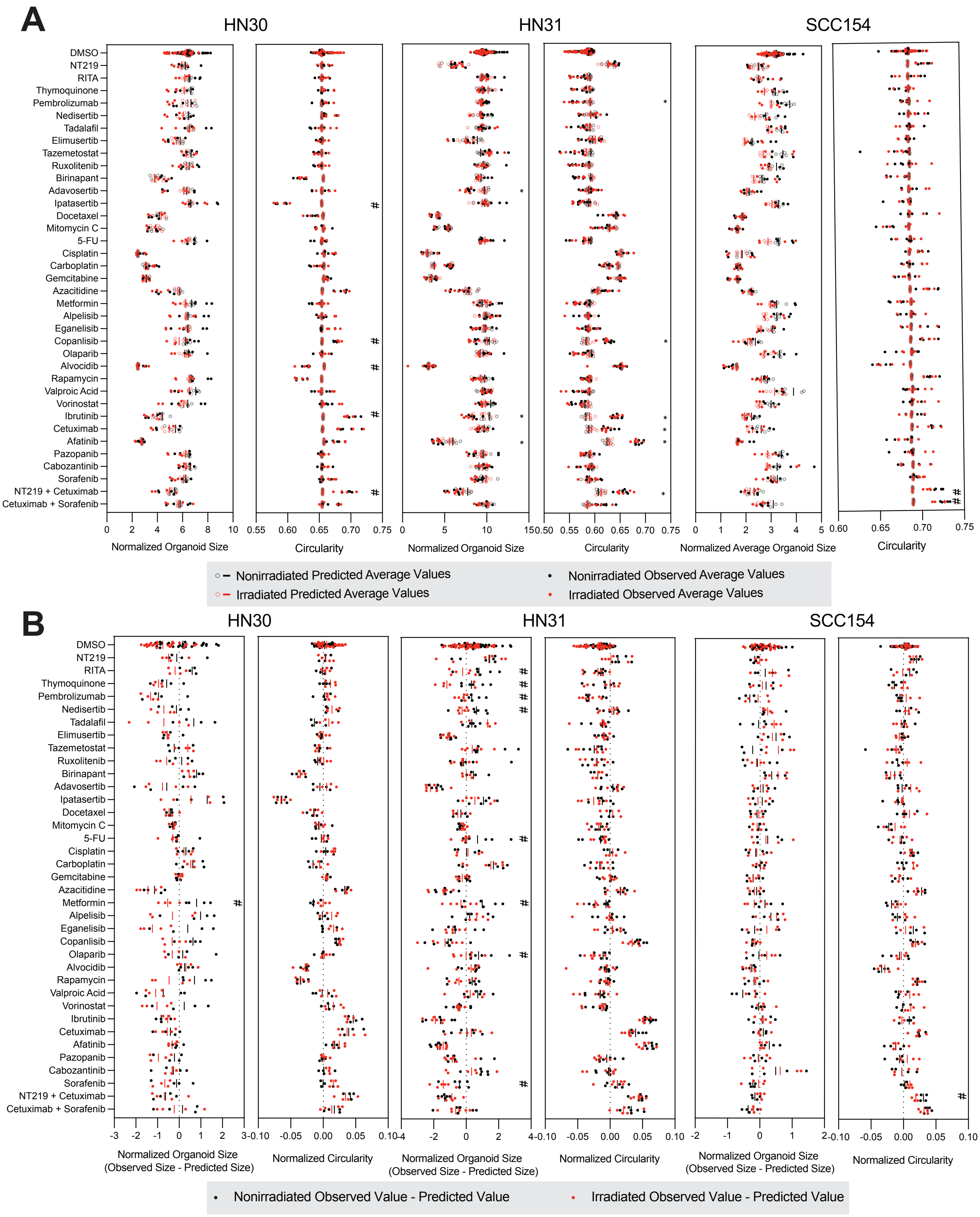
A
HN30
HN31
SCC154
DMSO
NT219
RITA
Thymoquinone
Pembrolizumab
Nedisertib
Tadalafil
Elimusertib
Tazemetostat
Ruxolitenib
Birinapant
Adavosertib
Ipatasertib
Docetaxel
Mitomycin C
5-FU
Cisplatin
Carboplatin
Gemcitabine
Azacitidine
Metformin
Alpelisib
Eganelisib
Copanlisib
Olaparib
Alvocidib
Rapamycin
Valproic Acid
Vorinostat
Ibrutinib
Cetuximab
Afatinib
Pazopanib
Cabozantinib
Sorafenib
NT219 + Cetuximab
Cetuximab + Sorafenib
Normalized Organoid Size
Circularity
Normalized Organoid Size
Circularity
Normalized Average Organoid Size
Circularity
Nonirradiated Predicted Average Values
Irradiated Predicted Average Values
Nonirradiated Observed Average Values
Irradiated Observed Average Values
B
HN30
HN31
SCC154
Normalized Organoid Size (Observed Size - Predicted Size)
Normalized Circularity
Normalized Organoid Size (Observed Size - Predicted Size)
Normalized Circularity
Normalized Organoid Size (Observed Size - Predicted Size)
Normalized Circularity
Nonirradiated Observed Value - Predicted Value
Irradiated Observed Value - Predicted Value

**Supplementary Figure 6.** Related to **Figure 2. Influence of Radiation onto Organoid Size and Circularity. (A)** Scatter plots showing observed and predicted average Day 9 organoid size and circularity for HN30, HN31, and SCC154 under various therapeutic agents, with or without radiation. Solid dots represent observed values per well measured with machine learning–based analysis of brightfield images. Translucent rings represent predicted values per well, calculated from normalized viability using linear regression models correlating viability with normalized organoid size or circularity without radiation. Bars denote mean predicted values for each treatment. Nonirradiated organoids are shown in black, and irradiated organoids in red. Student's t-test was used to compare predicted and observed values in nonirradiated organoids. *: $q < 0.01$; #: $0.01 < q < 0.02$. **(B)** Scatter plots showing the differences between observed and predicted average Day 9 organoid size and circularity for the same cell lines and treatments, with or without radiation. Dots represent the per-well difference between observed and predicted values. Bars denote mean differences for each treatment. Nonirradiated organoids are shown in black, and irradiated organoids in red. Student's t-test was used to compare differences between nonirradiated and irradiated conditions. #: $q < 0.35$.

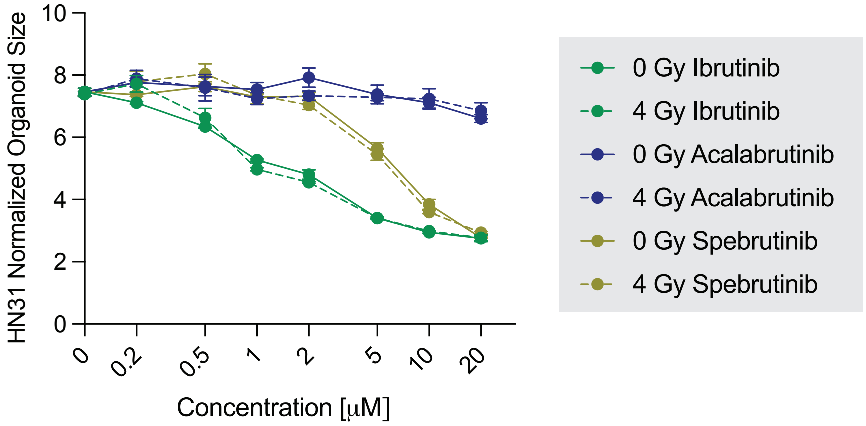


**Supplementary Figure 7.** Related to **Figure 3. HN31 BTK analysis Organoid Size Analysis.** Dose-response curves of Day 9 normalized average size of organoids of HN31 under treatment of Ibrutinib, Acalabrutinib or Spebrutinib of different concentrations. Organoid sizes are normalized to those of DMSO control organoids on Day 1. Data under treatment of Ibrutinib are shown in blue, data under Acalabrutinib blue, and data under Spebrutinib brown. Data of nonirradiated organoids are shown as solid lines, and data of irradiated ones dashed lines. Data are represented as mean ± SEM.

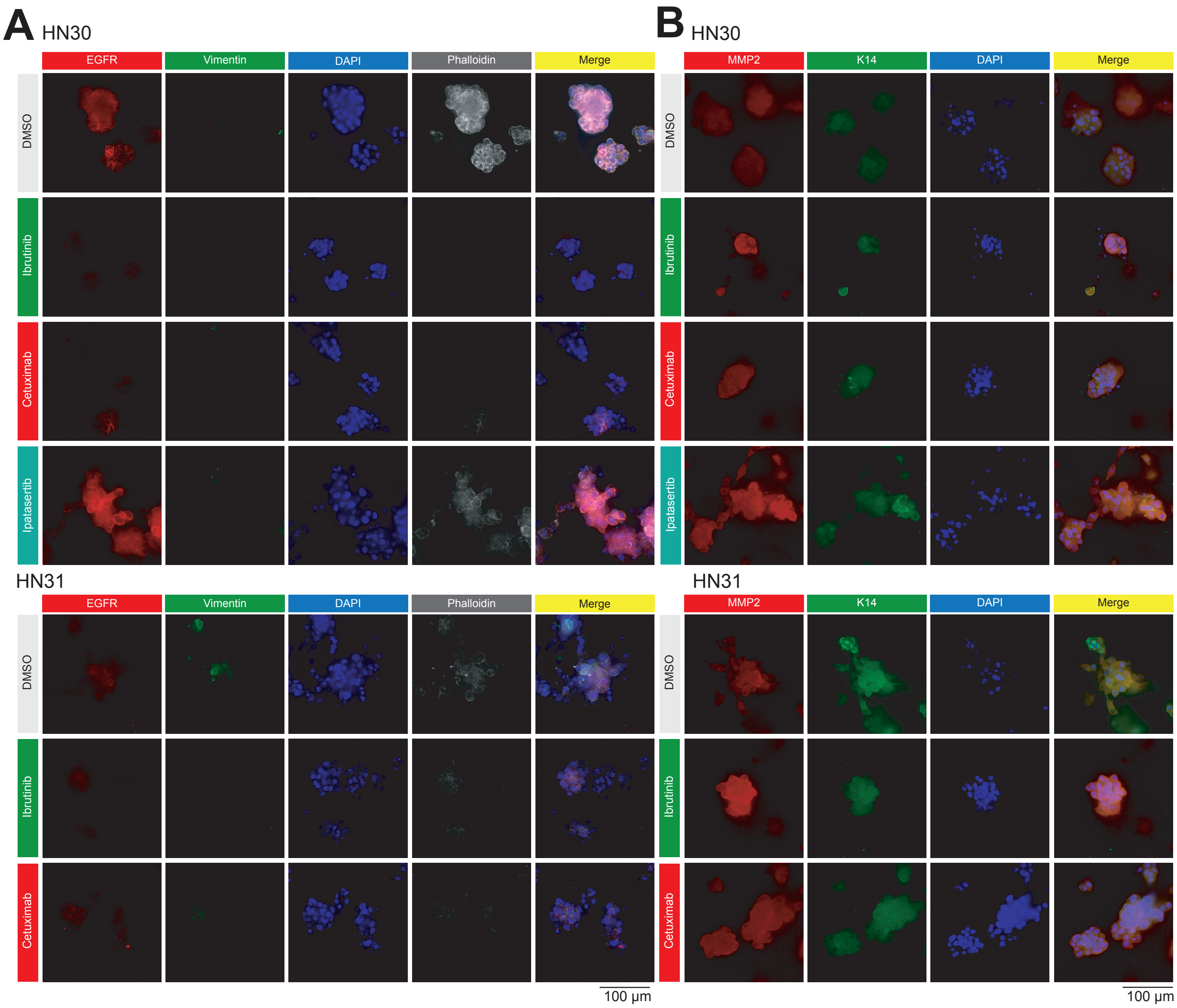


**Supplementary Figure 8.** Related to **Figure 4. Immunofluorescence Images of HN30 and HN31 Irradiated Organoids Stained with EGFR, Vimentin, Phalloidin, MMP2 and K14. (A)** Representative immunofluorescence images showing EGFR (red), vimentin (green), nuclei (DAPI, blue) and F-actin (Phalloidin, white) in irradiated HN30 and HN31 organoids treated with DMSO vehicle, Ibrutinib, Cetuximab or Ipatasertib (HN30 only). Scale bar: 100 µm for immunofluorescence images. **(B)** Representative immunofluorescence images showing MMP2 (red), K14 (green) and nuclei (DAPI, blue) in irradiated HN30 and HN31 organoids treated with DMSO vehicle, Ibrutinib, Cetuximab or Ipatasertib (HN30 only). Scale bar: 100 µm for immunofluorescence images.

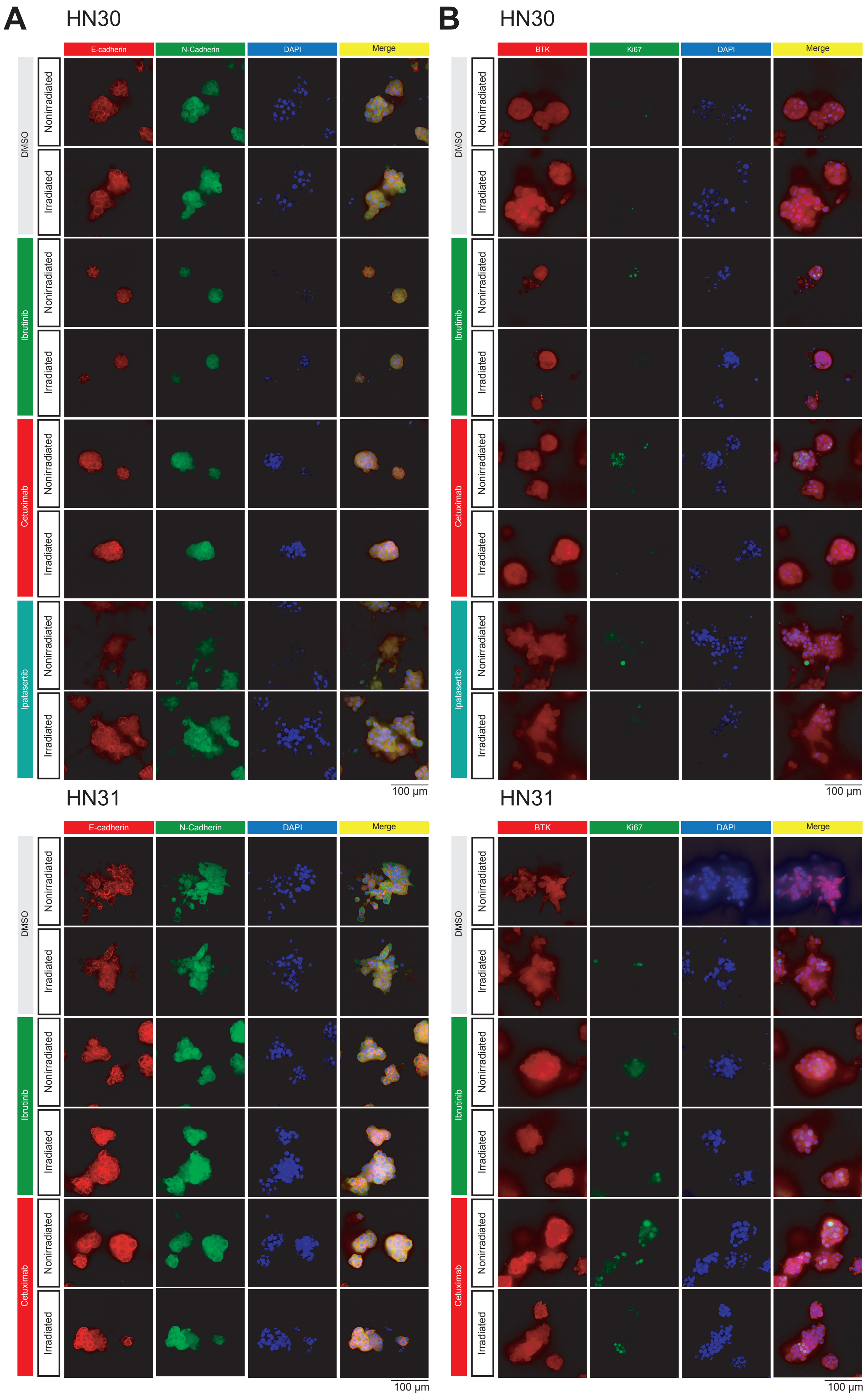
A
HN30
E-cadherin
N-Cadherin
DAPI
Merge
DMSO
Ibrutinib
Cetuximab
Ipatasertib
Nonirradiated
Irradiated
100 µm
HN31
B
BTK
Ki67

**Supplementary Figure 9.** Related to **Figure 4. Immunofluorescence Images of HN30 and HN31 Organoids Stained with E-cadherin, N-cadherin, BTK and Ki67. (A)** Representative immunofluorescence images showing E-cadherin (red), N-cadherin (green) and nuclei (DAPI, blue) in nonirradiated and irradiated HN30 and HN31 organoids treated with DMSO vehicle, Ibrutinib, Cetuximab or Ipatasertib (HN30 only). Scale bar: 100 µm for immunofluorescence images. **(B)** Representative immunofluorescence images showing BTK (red), Ki67 (green), nuclei (DAPI, blue) and F-actin (Phalloidin, white) in nonirradiated and irradiated HN30 and organoids treated with DMSO vehicle, Ibrutinib, Cetuximab or Ipatasertib. Scale bar: 100 µm for immunofluorescence images.

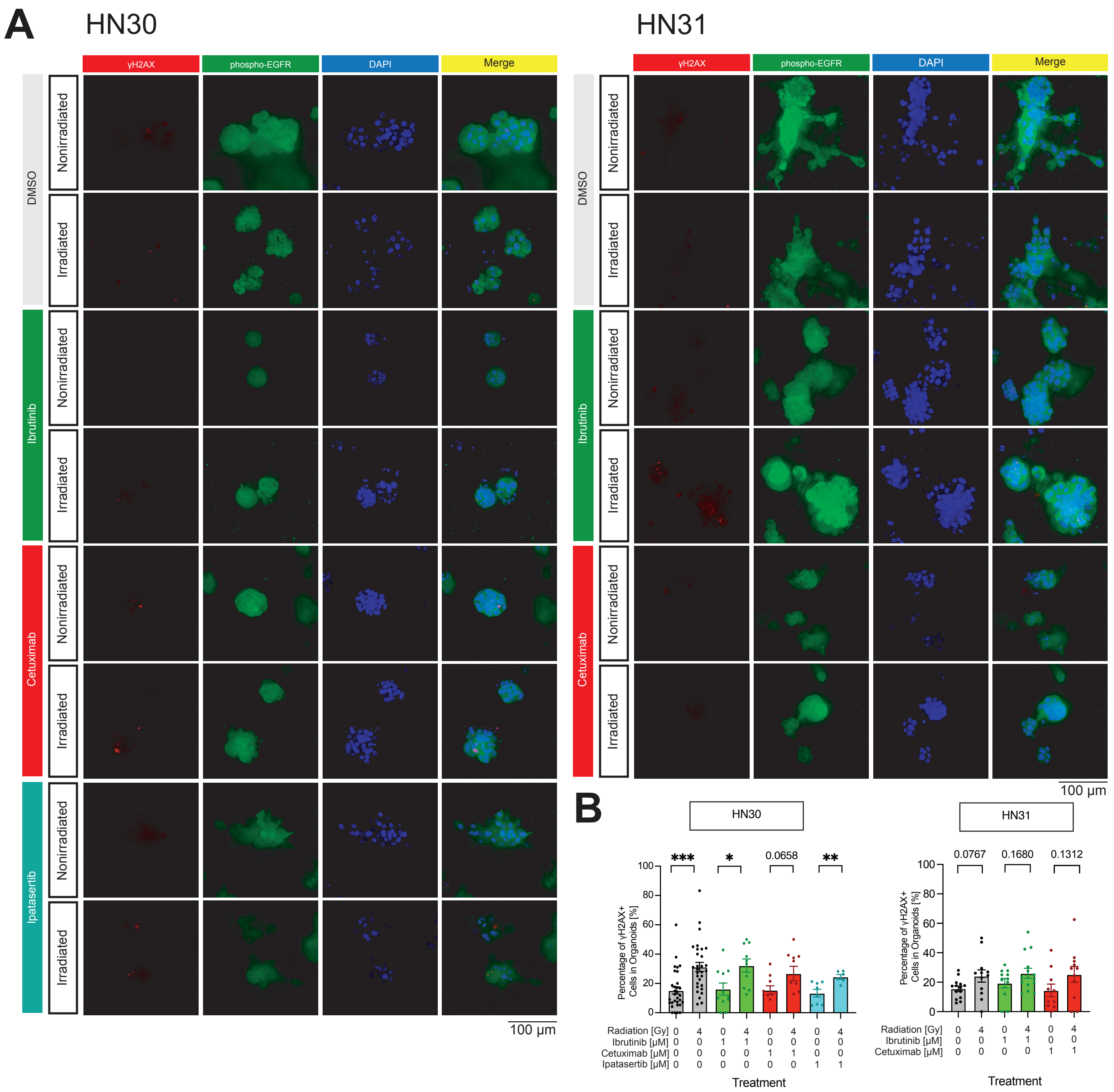


**Supplementary Figure 10.** Related to **Figure 4. Immunofluorescence Images of HN30 and HN31 Organoids Stained with γH2AX and Phospho-EGFR and γH2AX. (A)** Representative immunofluorescence images showing γH2AX (red), phospho-EGFR (green), nuclei (DAPI, blue) and F-actin (Phalloidin, white) in nonirradiated and irradiated HN30 and HN31 organoids treated with DMSO vehicle, Ibrutinib, Cetuximab or Ipatasertib (HN30 only). Scale bar: 100 µm for immunofluorescence images. **(B)** Bar plots of percentages of cells with γH2AX particles in each organoid of HN30 and HN31, with or without radiation, under treatment of Ibrutinib, Cetuixmab and Ipatasertib. Each dot represents the percentages of cells with γH2AX particles in each organoid, with total cells measured by counting DAPI particles. Student's t-test is done to compare the average percentages of irradiated and nonirradiated groups of each treatment. Data are represented as mean ± SEM. *: $0.0021 \leq$ p-value $< 0.0332$, **: $0.0002 \leq p < 0.0021$, ***: $0.0001 \leq p < 0.0002$:, ****: $p < 0.0001$.

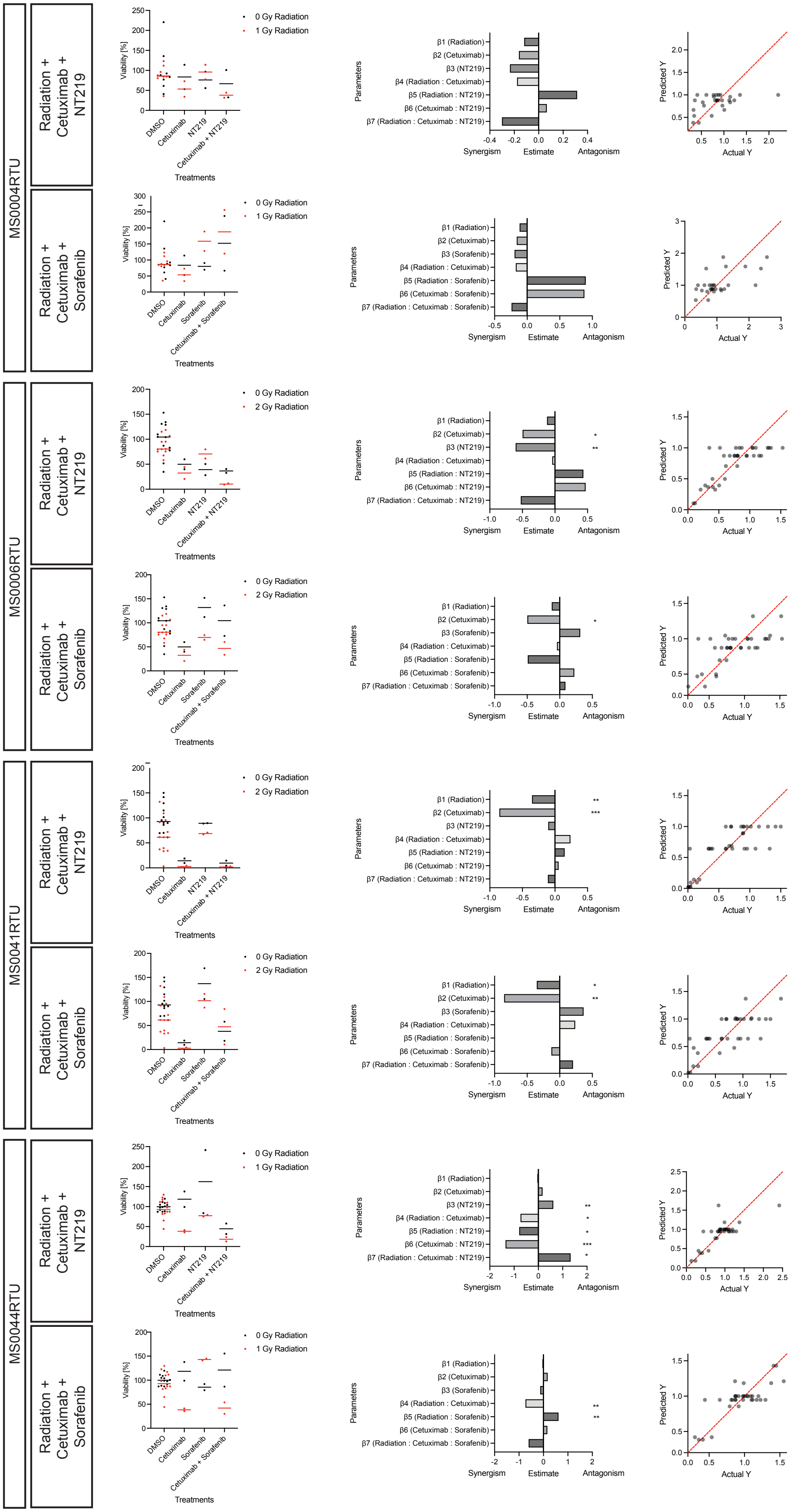
MS0004RTU
MS0006RTU
MS0041RTU
MS0044RTU
Radiation + Cetuximab + NT219
Radiation + Cetuximab + Sorafenib
Viability [%]
Treatments
DMSO
Cetuximab
NT219
Sorafenib
Cetuximab + NT219
Cetuximab + Sorafenib
0 Gy Radiation
1 Gy Radiation
2 Gy Radiation
Parameters
β1 (Radiation)
β2 (Cetuximab)
β3 (NT219)
β3 (Sorafenib)
β4 (Radiation : Cetuximab)
β5 (Radiation : NT219)
β5 (Radiation : Sorafenib)
β6 (Cetuximab : NT219)
β6 (Cetuximab : Sorafenib)
β7 (Radiation : Cetuximab : NT219)
β7 (Radiation : Cetuximab : Sorafenib)
Synergism
Estimate
Antagonism
Predicted Y
Actual Y

**Supplementary Figure 11.** Related to **Figure 6. Combination effects of Cetuximab and NT219 or Sorafenib in Clinical Samples. Left columns:** Normalized viability % of organoids of MS0004RTU, MS0006RTU, MS0041RTU and MS0044RTU treated with DMSO vehicles, Cetuximab, NT219 or Sorafinib, or combination with Cetuximab and NT219 or Sorafenib, with or without radiation. Data not under radiation are shown in black, and data under radiation are shown in red. **Middle columns:** Data of organoid viability of MS0004RTU, MS0006RTU, MS0041RTU and MS0044RTU under various treatments were analyzed with multiple linear regression, with a formula of Viability = β0 * Intercept + β1 * Radiation + β2 * Cetuximab + β3 * NT219 + β4 * Radiation: Cetuximab + β5 * Radiation : Agent + β6 * Cetuximab : Agent + β7 * Radiation : Cetuximab : Agent. Colons denote combination effects among two or three treatments. Agent is NT219 or Sorafenib. Bar plots of β1-6. *: 0.0021 ≤ p-value < 0.0332, **: 0.0002 ≤ p < 0.0021:, ***: 0.0001 ≤ p < 0.0002:, ****: p < 0.0001; **Right columns:** The scatter plot of the actual Y values (measured viability, X axis) and predicted Y values (predicted viability, Y axis). The more the data scatter along the line of identity (the red line), the more useful the model is.

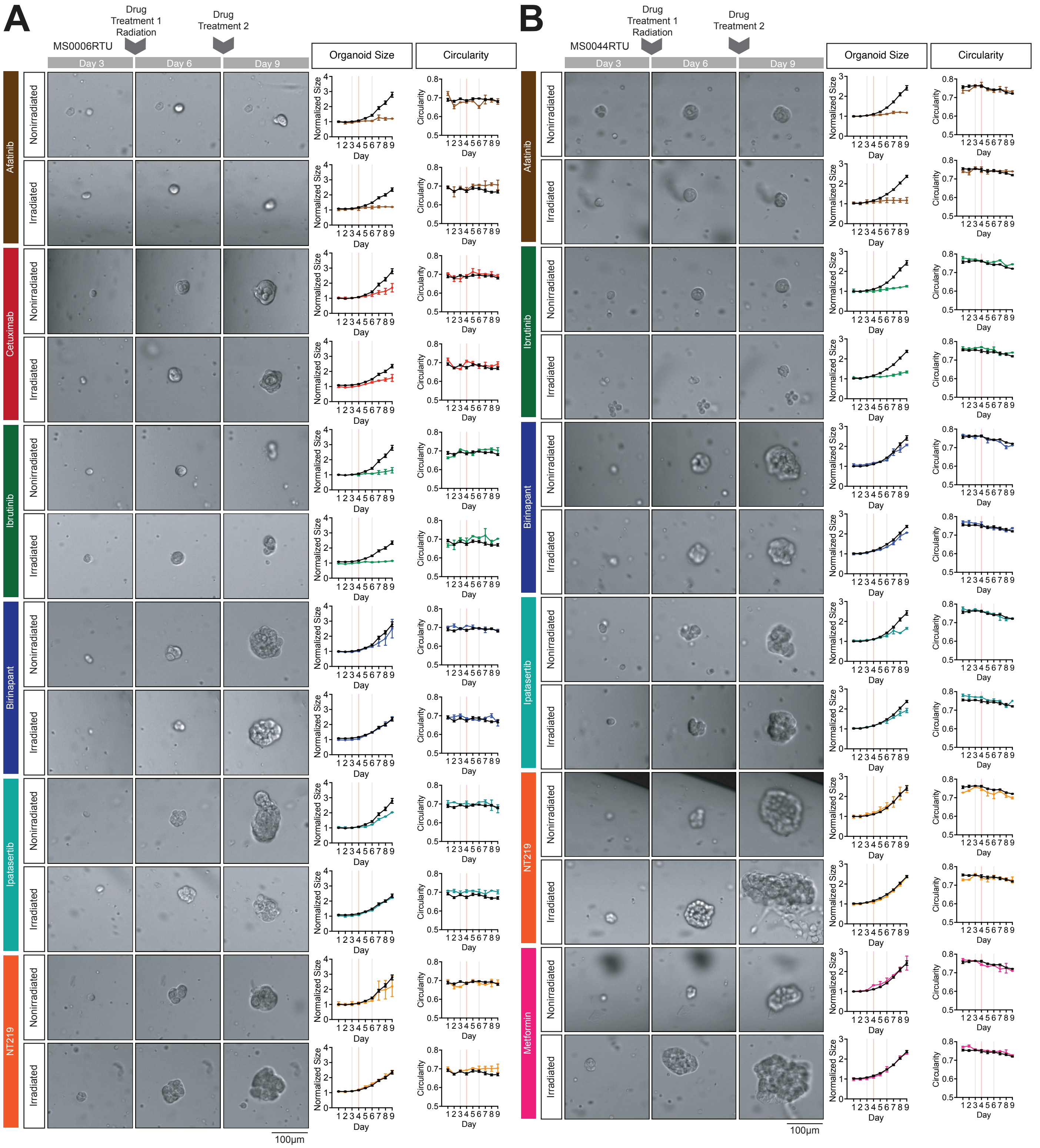


**Supplementary Figure 12.** Related to **Figure 7. Images of organoids and line charts of MS0006RTU and MS0044RTU. (A-B)** Supplementary representative brightfield images of organoids of MS0006RTU **(A)** and MS0044RTU **(B)** in culture on Day 3 (Column 1), Day 6 (Column 2) and Day 9 (Column 3). Organoids were treated with therapy agents on Day 3 and Day 6, and radiation on Day 4, after imaging. Growth (Column 4) and changes of morphology of organoids (Column 5) was tracked over time by segmenting in-focus organoids in the brightfield images using a machine learning–based pipeline. Growth was measured with the normalized organoid size every day, which is calculated by normalizing average cross-sectional area of all organoids across all wells of the same treatment of the same cell line across all trials to that of DMSO without radiation measured on Day 1 of culture. Data of control organoids are denoted with black lines. Data of treated organoids are denoted with respective colors. Data are represented as mean ± SEM. Morphology was measured with average circularity of all organoids across all wells of the same treatment of the same cell line across all trials. Scale bars: 100 μm for brightfield pictures.

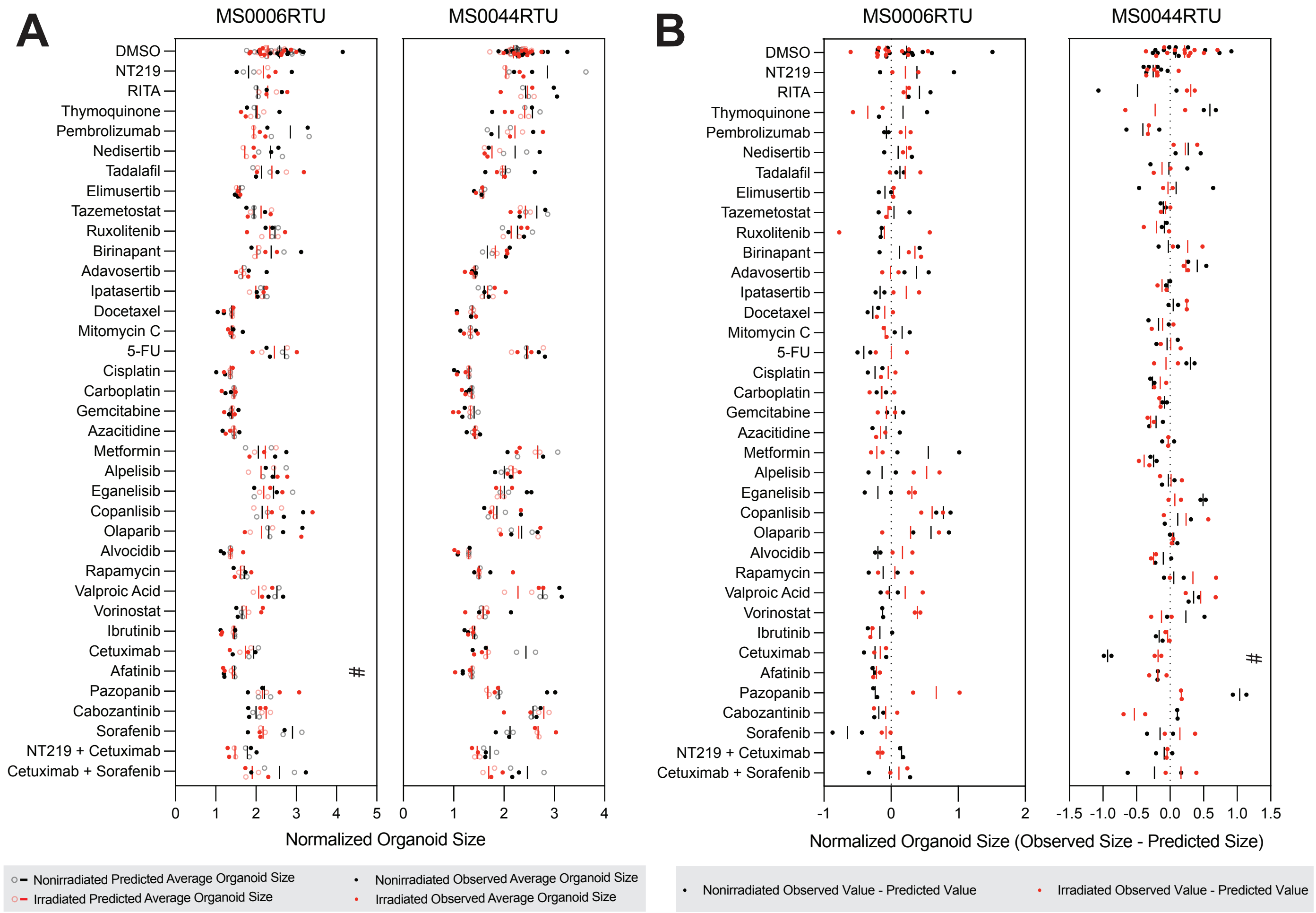


**Supplementary Figure 13.** Related to **Figure 7. Influence of Radiation onto Organoid Size in Clinical Samples. (A)** Scatter plots showing observed and predicted average Day 9 organoid size for MS0006RTU and MS0044RTU under various therapeutic agents, with or without radiation. Solid dots represent observed values per well measured with machine learning–based analysis of brightfield images. Translucent rings represent predicted values per well, calculated from normalized viability using linear regression models correlating viability with normalized organoid size or circularity without radiation. Bars denote mean predicted values for each treatment. Nonirradiated organoids are shown in black, and irradiated organoids in red. Statistical comparisons between predicted and observed values in nonirradiated organoids were performed using Student's t-test. For MS0006RTU, #: $q < 0.5$; for MS0044RTU, #: $q < 0.4$. **(B)** Scatter plots showing the differences between observed and predicted average Day 9 organoid size for the same clinical samples and treatments, with or without radiation. Dots represent the per-well difference between observed and predicted values. Bars denote mean differences for each treatment. Nonirradiated organoids are shown in black, and irradiated organoids in red. Student's t-test was used to compare differences between nonirradiated and irradiated conditions. For MS0044RTU, $q < 0.4$.
